# TUMOR ENDOTHELIAL CELL AUTOPHAGY IS A KEY VASCULAR-IMMUNE CHECKPOINT IN MELANOMA

**DOI:** 10.1101/2023.04.21.537799

**Authors:** Jelle Verhoeven, Kathryn A Jacobs, Francesca Rizzollo, Francesca Lodi, Yichao Hua, Joanna Poźniak, Adhithya Narayanan Srinivasan, Diede Houbaert, Gautam Shankar, Sanket More, Marco B Schaaf, Nikolina Dubroja Lakic, Maarten Ganne, Jochen Lamote, Johan Van Weyenbergh, Louis Boon, Oliver Bechter, Francesca Bosisio, Mathieu JM Bertrand, Jean Christophe Marine, Diether Lambrechts, Gabriele Bergers, Madhur Agrawal, Patrizia Agostinis

**Affiliations:** Cell Death Research and Therapy Laboratory, Center for Cancer Biology, VIB; Department of Cellular and Molecular Medicine, KU Leuven, Leuven, Belgium; Laboratory of Translational Genetics, Center for Cancer Biology, VIB; Department of Human Genetics, KU Leuven, Leuven, Belgium; Laboratory of Tumor Microenvironment and Therapeutic Resistance Center for Cancer Biology, VIB; Department of Oncology, KU Leuven, 3000 Leuven, Belgium; Laboratory of Translational Cell and Tissue Research, Department of Pathology, KULeuven and UZ Leuven, Belgium; Department of Pathology, UZLeuven, Belgium; Laboratory for Molecular Cancer Biology, Center for Cancer Biology, VIB; Laboratory of Clinical and Epidemiological Virology, Department of Microbiology, Immunology and Transplantation, Rega Institute for Medical Research, KU Leuven, Leuven, Belgium; Polpharma Biologics, Utrecht, the Netherlands; Department of General Medical Oncology UZ Leuven,Leuven; VIB Center for Inflammation Research, Ghent University; Department of Biomedical Molecular Biology, Ghent University, 9052 Ghent, Belgium

**Author notes:** Correspondence: Patrizia Agostinis, Madhur Agrawal. These authors contributed equally.

**Keywords:** autophagy, cancer, tumor endothelial cells, inflammation, immunotherapy

## Abstract

Tumor endothelial cells (TECs) actively repress inflammatory responses and maintain an immune-excluded tumor phenotype. However, the molecular mechanisms that sustain TEC-mediated immunosuppression remain largely elusive. Here, we show that autophagy ablation in TECs boosts antitumor immunity by supporting infiltration and effector function of T cells, thereby restricting melanoma growth. In melanoma-bearing mice, loss of TEC autophagy leads to the transcriptional expression of an immunostimulatory/inflammatory TEC phenotype driven by heightened NF-kB and STING signaling. In line, single-cell transcriptomic datasets from melanoma patients disclose an enriched Inflammatory^High^/Autophagy^Low^ TEC phenotype in correlation with clinical responses to immunotherapy. Congruently, patients responding to immunotherapy exhibit an increased presence of inflamed vessels, interfacing with infiltrating CD8+ T cells. Mechanistically, STING-dependent immunity in TECs is not critical for the immunomodulatory effects of autophagy ablation, since NF-kB-driven inflammation remains functional in STING/ATG5 double knockout TECs. Hence, autophagy is a principal tumor vascular anti-inflammatory mechanism dampening melanoma antitumor immunity.

## INTRODUCTION

The success of immunotherapy using immune checkpoint blockers (ICBs) has been remarkable in boosting CD8+ T cell-mediated antitumor immunity in the treatment of various solid tumors, including melanoma^1,2^. However, a substantial number of melanoma patients still fail to respond to immunotherapy^3^. Melanoma is a prototypical immunogenic tumor that can maintain a profound immunosuppressive tumor microenvironment (TME) by coopting multiple cancer cell-intrinsic and extrinsic mechanisms^2,4^. Together, these mechanisms limit the therapeutic efficacy of ICBs. Therefore, unravelling the molecular mechanisms that facilitate the immunosuppressive TME is a primary clinical need to overcome resistance to immunotherapy in melanoma.

Infiltration of CD8+ T cells into the tumor parenchyma is associated with a favorable prognosis and improved response to ICBs^2,5^. In particular, recent studies have indicated that the density, distribution, and activation pattern of tumor-infiltrating lymphocytes (TILs) in primary melanoma has prognostic significance^6^. In a recent phase 3 trial of patients with advanced melanoma, expanding the number of tumor-reactive T lymphocytes through autologous TIL-therapy, showed superior efficacy compared to anti-PD1 therapy^7^. This suggests that facilitating the infiltration of tumor reactive T lymphocytes within the tumor parenchyma may be critical in overcoming the immunosuppressive TME in melanoma.

In order to recognize cancer cells and eliminate them, cytotoxic CD8+ T cells need to extravasate the vascular lumen, transmigrate the endothelial lining to infiltrate the tumor, survive within the TME and evade immunosuppressive signals. In advanced tumors, vessel remodeling through aberrant angiogenesis generates a barrier to T cell infiltration by heightening the immunosuppressive features of TECs. TECs impair lymphocyte recruitment and transmigration (diapedesis) by reducing the secretion of immunoattracting chemokines (e.g.CXCL10, CXCL1), expression of adhesion molecules (e.g. vascular cell adhesion protein 1 (VCAM1), intercellular adhesion molecule 1 (ICAM1), E-and P-selectins) and the antigen presentation machinery (e.g. MHC class I) on their surface^8–10^. Through these immunosuppressive processes TECs impair antitumor immunity^11,12^ and represent a bottleneck for immunotherapies. Unraveling key mechanisms capable of counteracting the immunosuppressive features of TECs, or endorsing their inflamed status, could be crucial to fully unleash antitumor immune responses.

Macroautophagy (hereafter called autophagy) is a vital regulator of the immune and inflammatory microenvironment in tumors and other inflammatory conditions^13^. Autophagy is the primary lysosomal degradation pathway responsible for the degradation of intracellular material to maintain cell survival and energy homeostasis^14^. In advanced tumors, autophagy supports the heightened metabolic needs of cancer cells, and exerts cell non-autonomous functions through the unconventional secretion of inflammatory mediators^15,16^. Recent studies have indicated that loss of autophagy in the host tissues induces systemic and tissue-specific metabolic reprogramming that favors antitumor immunity. Full-body or liver-specific deletion of the canonical autophagy genes *Atg5* or *Atg7* impairs tumor immunotolerance and the growth of tumors with high mutational burdens, while remaining ineffective against tumors with a low mutational load^17^. T cells from *Atg5*^-/-^ mice display greater T cell-intrinsic effector function resulting from enhanced glycolytic metabolism and transcriptional upregulation of both metabolic and effector target genes^18^.

While these studies support the immunosuppressive nature of host autophagy, it is not well understood how autophagy regulates inflammatory/immunomodulatory functions of blood vessels at the interface with immune cells, under physiological or pathological conditions. Recently, in a model of acute physiological inflammation, genetic loss of autophagy in venular ECs of cremaster muscles exacerbated neutrophil transendothelial migration and tissue damage in a model of acute physiological inflammation, by controlling degradation of adhesion molecules at the sites of EC contacts^19^. Yet, it remains unknown to which extent TEC-associated autophagy shapes inflammation, immunosurveillance and responses to ICBs.

In this study, we combined multiplexed analysis and single-cell transcriptomic profiling of TECs from human melanoma, with *in vivo* functional/mechanistic studies in melanoma-bearing mice in which different autophagy genes were conditionally deleted in ECs. We demonstrate that autophagy in TECs exerts specific and persuasive anti-inflammatory and immunosuppressive functions, which impede antitumor immunity and responses to immunotherpy.

## RESULTS

### LOSS OF AUTOPHAGY IN ENDOTHELIAL CELLS IMPROVES MELANOMA IMMUNOSURVEILLANCE

To uncover the contribution of EC-autophagy in regulating immunosurveillance in melanoma, we crossed the inducible *PDGFb-cre^ERT^*^2^ *Rosa26^tdTomato/tdTomato^ line* with *Atg5*^fl/fl^ mice to delete the essential autophagy gene *Atg5* from blood ECs (i.e. Atg5^BECKO^) upon tamoxifen administration. Loss of exon 3 of *Atg5* was verified using qPCR on sorted murine TECs from B16-F10 melanoma-bearing mice and by change in reporter protein expression (switch from dTomato to GFP after deletion) by flow cytometry and microscopy (Extended Data Fig. 1a-c). Genetic loss of *Atg5* in TECs increased the presence of p62 puncta in CD31 positive vessels, indicating blockade in autophagy flux (Extended Data Fig. 1d), whereas vessel maturation as measured by aSMA+ and NG2+ pericyte coverage of CD31+ TECs was not significantly changed (Extended Data Fig. 1e,f).

We then compared the growth of subcutaneously implanted murine B16-F10 melanoma, a prototypical poorly immunogenic tumor, or the more immunogenic HCmel12 (mCherry+) melanoma, derived from a serial transplant of a UV-induced primary melanoma in *Hgf-Cdk4^R24C^* mice ^20^, in WT and Atg5^BECKO^ mice. In both syngeneic melanoma models, deletion of *Atg5* in blood ECs yielded a similar reduction in tumor burden (Fig. 1a-d). To exclude the possibility that these differences were caused by possible non-autophagic functions of Atg5^21^, we included Atg12^ECKO^ or Atg9^ECKO^ mice which were generated by crossing *Atg12^f^*^l/fl^ or *Atg9^f^*^l/fl^ mice with the *VeCadh-cre^ERT^*^2^ line^22^. As the canonical autophagy Atg5 and Atg12 proteins are part of the same multimeric ubiquitin-like conjugation complex that regulates the developing autophagosome^23^, we reasoned that *Atg*12 deletion should be functional epistatic to the loss of *Atg5* in TECs. Atg9 is a transmembrane protein that is phosphorylated by ULK1 and recruited to the PI3KC3 autophagy initiation machinery complex, which governs the first steps of autophagosome formation^24^. Moreover, while both Atg5 and Atg12 are also involved in LC3-associated phagocytosis, components of the ULK1 complex including Atg9 are not^21^. We observed similar phenotypic differences in tumor burden when comparing WT and Atg12^ECKO^ or Atg9^ECKO^ mice inoculated with the immunogenic YUMMER 1.7 melanoma cells, carrying the *Braf*^V600E^,*Pten*^−/−^/*Cdkn2a*^−/−^ mutations, or B16-F10 melanoma, respectively (Fig. 1e-h). In both B16-F10 and YUMMER 1.7 tumors, loss of TEC-autophagy increased the presence of CD3+ T lymphocytes around CD31+ TECs, particularly at the more vascularized tumor edge (Fig. 1i-j), suggesting an increased ability to attract and recruit T cells, irrespective of the immunogenicity of the melanoma models used.

**Figure 1:**
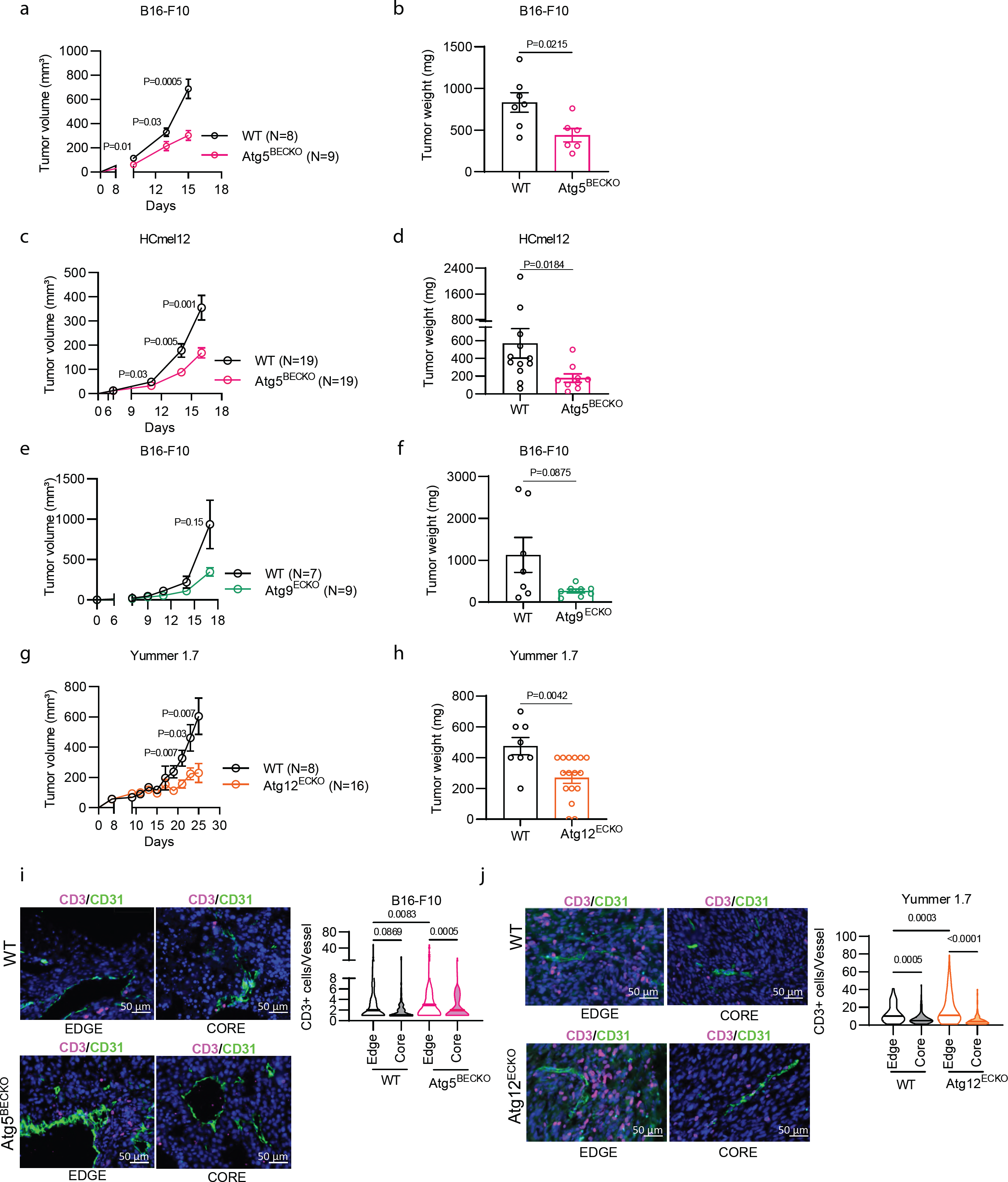
Genetic loss of autophagy in endothelial cells decreases growth of melanoma. **a-h,** Tumor volume **(a,c,e,g)** and end-point tumor weights **(b,d,f,h)** of WT and Atg5^BECKO^ mice subcutaneously injected with B16-F10 **(a,b)** and HcMel12-mcherry+ **(c,d)** melanoma cells; WT and Atg9a^ECKO^ mice subcutaneously injected with B16-F10 melanoma **(e,f)**; WT and Atg12^ECKO^ mice subcutaneously injected with Yummer 1.7 melanoma cells **(g,h)**. **i,j,** Representative images and quantification of immunofluorescence staining for CD3^+^ T cells in the edge and the rim of tumor sections of subcutaneous B16-F10 from WT and Atg5^BECKO^ mice **(i)** or Yummer 1.7 **(j)** tumors from WT and Atg12^BECKO^ mice. Scale bar represents 50µm. For immunofluorescence staining, at least 3 WT and 3 Atg5^BECKO^ mice and 2 WT and 3 Atg12^ECKO^ mice were used for the quantification. All data represent mean ± SEM. Statistical differences were determined using a two-sided Student’s t-test **(a-h)** or one way Anova with Holm-Sidak correction for multiple comparisons **(i,j)**. Source data are available for **a-j**.

We then analyzed TILs from B16-F10 bearing WT and Atg5^BECKO^ mice for the presence of CD8+ and CD4+ T cells by flow cytometry. Melanomas of Atg5^BECKO^ mice, compared to those of WT mice displayed an increased amount of both CD8+ T and CD4+ T cells (Fig. 2a,b) and harbored CD8+ T cells with higher expression of the key effector molecule Granzyme B (GrzB) (Fig. 2a). Melanoma-bearing Atg5^BECKO^ mice also exhibited a decreased ratio of immunosuppressive T regulatory cells (Tregs) to CD8+ T cells, an indicator of a diminished immunosuppressive status of the TME (Fig. 2c).

**Figure 2:**
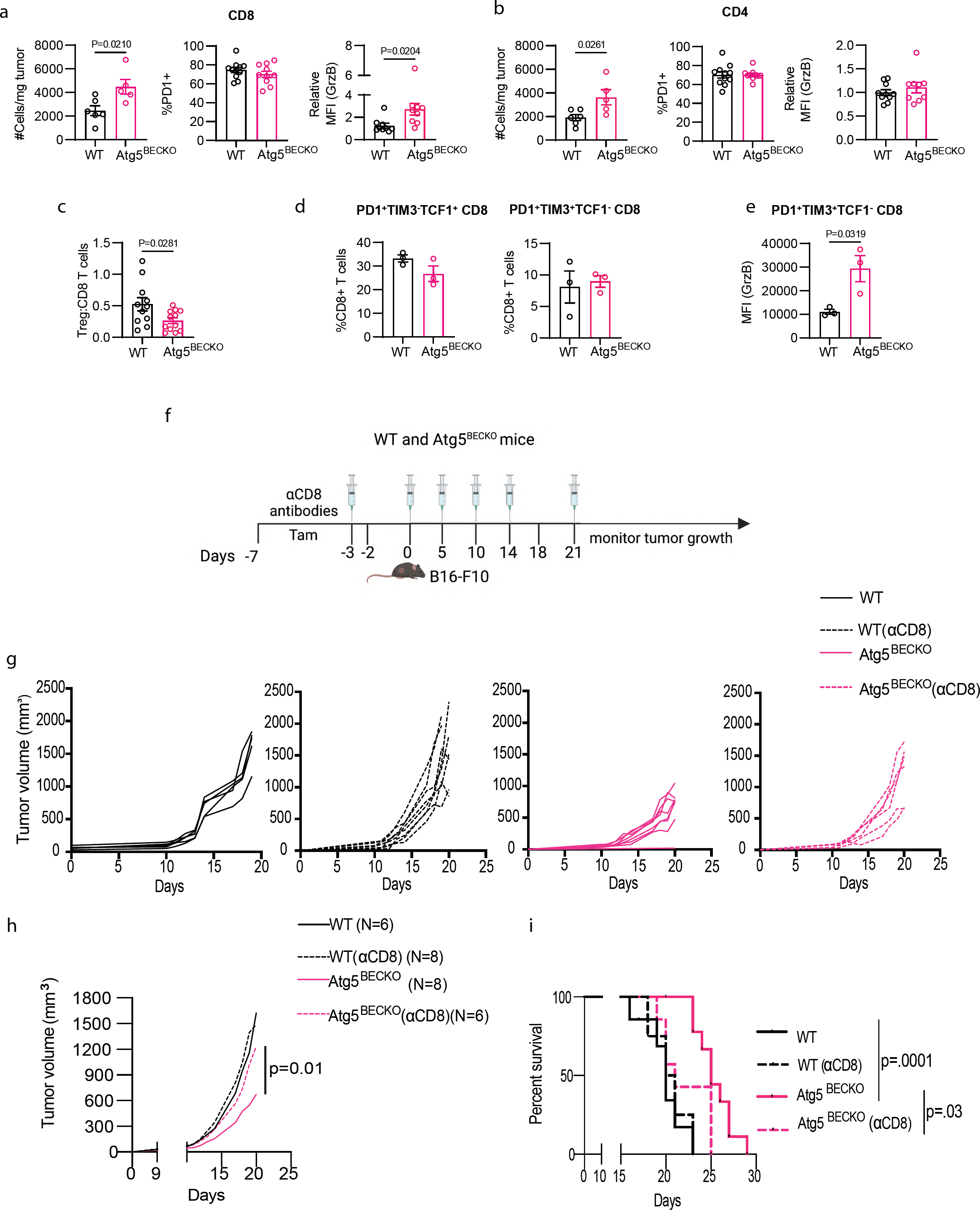
Deletion of *Atg5* in tumor endothelial cells promote CD8^+^ T cell mediated antitumor immune responses. **a-g,** Flow cytometry analysis of tumor infiltrating lymphocytes in subcutaneous B16-F10 tumors from WT and Atg5^BECKO^ mice. **a,** CD8^+^ T cells (# cells/mg tumor), PD1^+^ cells (% of CD8^+^ T cells) and Granzyme B (GrzB) (Mean fluorescence intensity or MFI in CD8^+^ T cells). **b,** CD4^+^ T cells (# cells/mg tumor), PD1^+^ cells (% of CD4^+^ T cells) and Granzyme B (GrzB) (MFI in CD4^+^ T cells). **c,** Ratio of # of CD4^+^FOXP3^+^ Tregs to CD8^+^ T cells. Data shown are representative of two independent experiments **(a-c)**. **d,** PD1^+^TIM3^+^TCF1^-^ cells (% of CD8^+^ T cells) and PD1^+^TIM3^-^TCF1^+^ cells (% of CD8^+^ T cells). **e,** intracellular Granzyme B (GrzB) of PD1^+^TIM3^+^TCF1^-^ cells. **f,** Schematic representation for depleting CD8^+^ cells in subcutaneous B16-F10 tumor bearing WT and Atg5^BECKO^ mice. **g,h,i**, Individual tumor volume **(g)** and end-point tumor volume **(h)** and survival analysis **(i)** of B16-F10 tumor bearing WT and Atg5^BECKO^ mice injected with vehicle or αCD8 antibody (100μg per mouse). All data represent mean ± SEM. Statistical differences were determined by two-sided Student’s t-test **(a-h)** or Kapelin-meyer survival analysis (i). Source data are available for **a-j**.

To gain further insight into the activated TIL populations, we profiled precursor exhausted CD8+ T cells (or T_PEX_) hallmarked by the combined expression of TCF1 (a stem-/memory-like marker) and PD1, and lack of TIM3 expression (as a terminally differentiated marker) and their progeny pool of terminally exhausted effector PD1+ CD8+ T cells (or T_EX_), defined by increased GrzB and high TIM3 levels, but lack of TCF1 expression (Extended data Fig. 2a) ^30^. Both populations remained similar in WT and Atg5^BECKO^ mice (Fig. 2d). However, terminally exhausted effector CD8+ T from Atg5^BECKO^ produced higher amounts of GrzB as compared to WT mice (Fig. 2e).

We then asked whether the increase in CD8+ T cells was causally responsible for the reduced tumor growth of the Atg5^BECKO^ mice mice. We depleted CD8+ T cells by injecting αCD8+ antibodies and examined tumor growth in WT and Atg5^BECKO^ mice (Fig. 2f). Depletion of CD8+ T cells (Extended data Fig. 2b) did not significantly affect B16-F10 tumor growth in WT mice, but reversed the beneficial effect on tumor burden and survival in Atg5^BECKO^ mice (Fig. 2g-i), thus functionally implicating CD8+ T cells in the delayed tumor growth phenotype of the Atg5^BECKO^ mice.

Together, these data suggest that loss of canonical autophagy genes in TECs restrains melanoma growth and the immunosuppressive TME by increasing the frequency of GrzB-expressing effector CD8+ T cells in the tumor.

### AUTOPHAGY FOSTERS AN IMMUNOTOLERANT STATUS OF TUMOR ENDOTHELIAL CELLS

Next, we asked by which means autophagy regulates lymphocyte influx and activity, by first focusing on lymphocyte-regulating chemokines, cytokines, and adhesion molecules. As autophagy not only exerts direct effects via protein degradation, but also regulates the transcriptional and epigenetic programs in cells, including ECs^22,25,26^, we initially conducted transcriptional profiling of flow-cytometry-sorted (FACS) ATG5-proficent and -deficient CD45^-^Ter119^-^CD31^+^ TECs from B16-F10 tumors (yielding more than 90% pure TECs), with a panel of 770 genes implicated in cancer immunomodulation using the Nanostring technology (Fig. 3a). The top hallmarks in our data set were related to inflammatory responses (Fig.3b). Moreover, analysis of differentially expressed (DE) genes with Log_2_FC>=0.7 and P<0.05, revealed a prominent expression of genes in Atg5^BECKO^ TECs encoding for surface inflammatory proteins with immune cell-interacting activities and cytokines/chemokines, which are characteristics of an inflamed TEC phenotype (Fig. 3b). Congruently, gene expression of factors implicated in the NF-κB pathway and various EC adhesion molecules were significantly upregulated in TECs from Atg5^BECKO^ mice. The latter included *VCAM1, ICAM1*, *SELE* and *SELP* with an established role in binding T cell surface ligands and in increasing T cell receptor signaling^27–29^, as well as cytokines and chemokines with chemoattractant and adhesion function in ECs such as *IL6, CX3CL1/fractalkine*, *CXCL2*, *CXCL1*, and *RELB* a mediator of the non canonical NF-κB pathway (Fig. 3c). Another set of DE genes annotated to the antiviral/type 1 interferon (IFN) responses and included *IFIH1,IRF1,IRF7,TLR4,* and elements of the antigen presentation machinery *H2-K1, H2-T23* and *H2-D1,* suggesting that TECs from Atg5^BECKO^ mice have acquired improved immunomodulatory functions, including the capacity to sustain T cell function (Fig. 3c). Gene set enrichment analysis (GSEA) confirmed that TECs from Atg5^BECKO^ mice compared to TECs from WT mice, exhibited a robust enrichment in gene sets involved in inflammatory response and interferon signaling pathways (Fig. 3b). Thus, deletion of Atg5 in TECs evokes the expression of lymphocyte attracting chemokines and molecules controlling lymphocyte adhesion and function.

**Figure 3:**
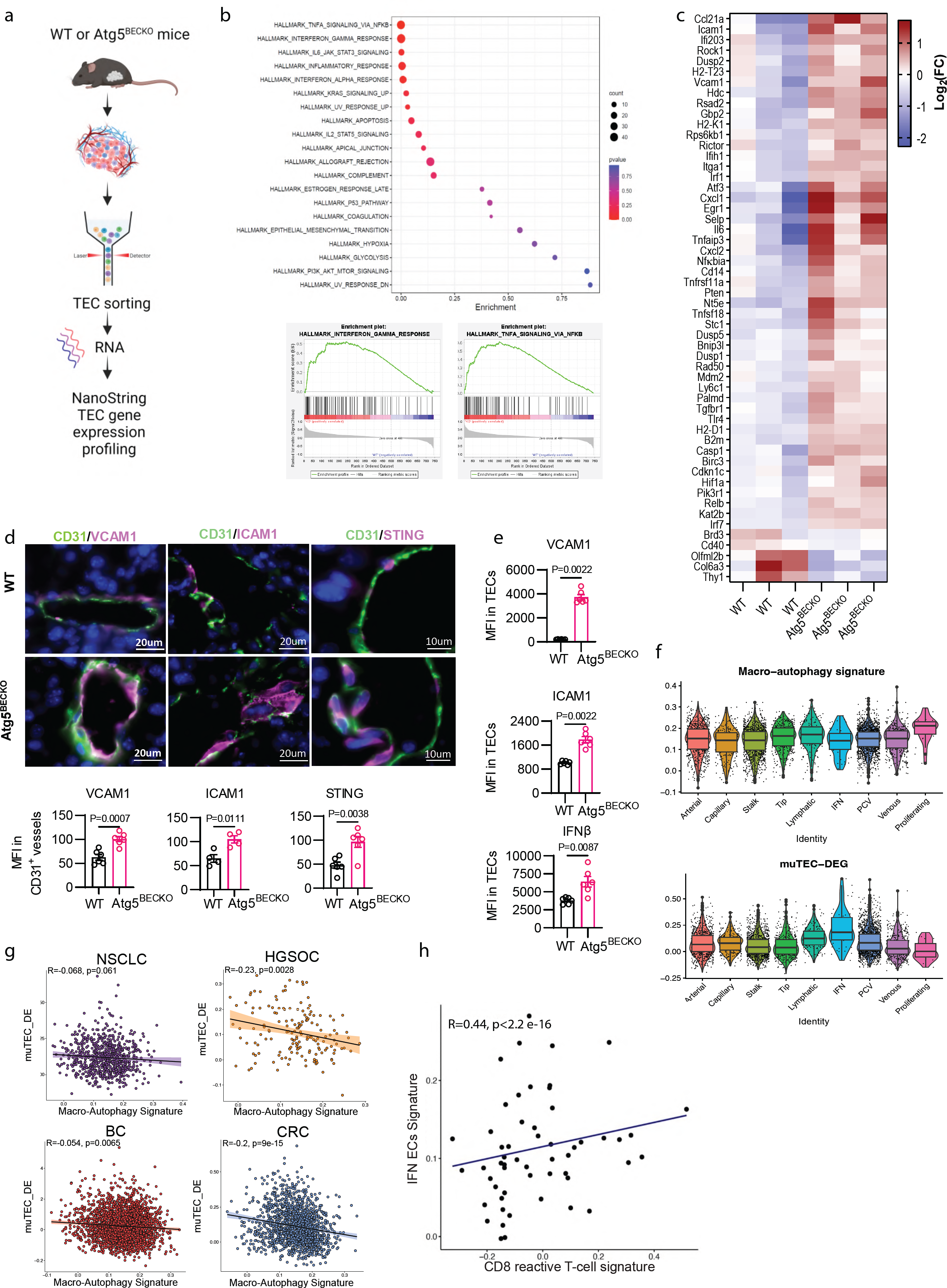
Autophagy represses inflammatory phenotype of Tumor Endothelial Cells. **a,** Experimental design for end-point investigation of tumor derived ECs. **b,c,** NanoString analysis from TECs in subcutaneous B16-F10 tumors from WT and Atg5^BECKO^ mice. **b,** Gene Set Enrichment Analysis (GSEA) analysis. **c,**Heap map showing differentially expressed genes (Log_2_FC >= 0.7, P<0.05). **d,** Representative images and quantification (MFI) of immunofluorescence staining for VCAM1, ICAM1 and STING in CD31+ tumor endothelial cells from tumor sections of subcutaneous B16-F10 tumors from WT and Atg5^BECKO^ mice. Scale bar represents 20µm. **e,** flow cytometry analysis of surface expression of VCAM1, ICAM1 and intracellular IFNβ in TECs in subcutaneous B16-F10 tumors from WT and Atg5^BECKO^ mice. **f,** Expression of macro-autophagy signature and muTEC-DEG across the 9 ECs subtypes (n ECs =4.921). Median of the data is displayed as solid black lines. **g**, Spearman correlation between the expression of macro-autophagy signature and mu-TEC-DE in ECs of primary tumours. **h**, Spearman correlation between the expression of CD8+ reactive T-cells signature (calculated on T-cells) and IFN ECs signature (calculated on ECs) in each primary tumor sample.For immunofluorescence staining, at least 3 WT and 3 Atg5^BECKO^ mice were used for the quantification. All data represent mean ± SEM. Statistical differences were determined by two-sided Student’s t-test (b-e) or **f-i**, Exact P value calculated by two-sided Mann-Whitney test or two-sided Wilcoxon matched-pairs signed rank test. Only primary tumor samples of HGSOC, CRC, BC and NSCLC are included in the analysis. Source data are available for **a-e**.

We then validated the transcriptome results by immunostaining for selected immunomodulatory /inflammatory markers. Compared to TECs of WT B16-F10 melanoma, TECs from from Atg5^BECKO^ mice demonstrated a significant increase in the levels of VCAM1, ICAM1 and of STING, a major regulator of the the antiviral/type 1 interferon responses^30^ (Fig. 3d).

Congruently, the surface expression of VCAM1, ICAM1 (Fig. 3e), MHC class I and MHC class II (Extended data Fig. 3a) and the intracellular expression of IFNb -a downstream effector of the STING pathway^30^-(Fig. 3e), were elevated in TECs from Atg5^BECKO^ mice. A similar inflammed TEC phenotype was also observed in YUMMER 1.7 melanoma grown in Atg12^ECKO^ mice (Extended Data Fig.3b). Thus, genetic blockade of canonical autophagy boosts TEC inflammation in various melanoma models.

We next sought to more broadly contextualize our murine mRNA profiling data to human TECs (huTECs), by analyzing phenotypic characteristics of ECs from publicly available single-cell RNA-seq (scRNA-seq) cancer atlases. To increase coverage of rare huTECs subtypes, we pooled EC scRNAseq data from non-small cell lung cancer (NSCLC), breast cancer (BC), high-grade serous ovarian carcinoma (HGSOC) and colorectal cancer (CRC), from different biopsy sites (primary tumors, metastasis and adjacent non-neoplastic tissue samples). Starting from a total of a 7.573 high quality TECs, we performed cluster analysis across these different tumors (Extended data Fig. 3c-f). HuTECs were clustered into nine distinct subtypes and validated using markers previously identified in human and murine ECs^11^ (Extended data Fig. 3c-e). These clusters included arterial (*MGP, FBLN5, GJA4*) and capillary ECs (*FABP4, CD36, CA4*), stalk cells *(INSR, COL4A1, FLT1*), tip cells (*PGF, TP53I11, ESM1*) and lymphatic ECs (*PROX1, PDPN, LYVE1*), as well as ECs characterized by an IFN response signature (*ISG15, CCL2, CXCL9*), suggesting their involvement in immune cell recruitment (Extended data Fig. 3c,d). Additionally, we distinguished venous from post-capillary venules (PCV), the latter being the most abundant ECs cluster (Extended data Fig. 3e) and found a minor subcluster of proliferating ECs (Proliferating; Extended data Fig 3e). Finally, we observed a variation of different huTECs subtypes frequency across tumor types, for example the IFN huTECs were predominantly present in NSCLC (Extended data Fig. 3f). We validated that the ‘inflammatory’ gene signature derived from the Nanostring analysis contained genes conserved between mouse and human (Extended Data Table 1), and used this in-house generated signature, dubbed as muTEC-DE, and the Reactome-macroautophagy signature (including *ATG5*, *ATG12* and *ATG9* among various genes involved in canonical autophagy)^31^ for further analysis. Notably, the IFN huTEC subset of primary tumors, hallmarked by the highest expression of the muTEC-DE signature, had the lowest enrichement in the autophagy signature (Fig. 3f), suggesting that, particularly the huTEC subtype with more pronounced immunomodulatory function, maintains low expression level of autophagy genes.

We then further tested the association between the muTEC-DE and autophagy signatures within huTECs of primary tumors, across cancer types. Remarkably, we observed a significant inverse correlations between the muTEC-DE and autophagy gene signatures in all of these cancer types (Fig 3g). Furthermore, enrichement in the muTEC-DE signature in huTECs showed a positive correlation with the expression of a CD8 + reactive T cells signature, measured within T cells (Fig. 3h).

Together these observations suggest that irrespective of the species (mouse or human) or tumor types, TECs with a low autophagic capacity have enhanced immunosupporting functions.

### AUTOPHAGY BLUNTS THE NF-κB AND cGAS-STING INFLAMMATORY AXIS IN ECs

We next set out to identify the underlying mechanisms by which inhibition of autophagy endorses the EC inflamed phenotype. We first evaluated whether cultured human umbilical ECs (HUVECs) could reliably model the effects of genetic loss of autophagy observed in mice. Deletion of *Atg5* by CRISPR-Cas9 in ECs blunted the conversion of LC3BI to LC3BII, and led to the accumulation of p62, indicating blockade of constitutive autophagic flux (Extended data Fig. 4a-c). No sign of caspase activation was observed under baseline/replete conditions (Extended data Fig. 4d) as also observed in our previous study^22^. Compared to control ECs (Ctr), Atg5KO ECs displayed enhanced expression of a largely similar set of inflammatory genes (e.g. *SELE, VCAM1, ICAM1)* and chemokines *(e.g., CX3CL1, CXCL10)* as observed in the Nanostring analysis of muTECs isolated from Atg5^BECKO^ mice (Fig. 4a-e). Similar results were obtained by inhibiting ULK1/2 kinase activity or by increasing the lysosomal pH through bafilomycin A (Extended data Fig. 4e), suggesting that under baseline conditions, inflammation in ECs is downregulated at the transcriptional level via the canonical lysosomal-autophagy pathway. Differences in the extent of inflammatory gene expression profiles (Fig. 4a-e, Extended data Fig. 4e) are likely explained by the acute chemical inhibition of ULK1/2 versus the chronic deletion of *Atg5* in these cells. Increased surface expression of VCAM1 in Atg5KO cells was further validated by Flow Cytometry (Fig. 4f, Extended data Fig. 4f). Therefore, to gain more functional evidence, we then tested the EC adhering ability of αCD3/CD28-activated JURKAT cells in co-culture with WT or Atg5 KO ECs. In this setting cytokines (i.e. IFNγ/TNFα) were supplemented to the EC cell culture medium to further promote surface translocation of adhesion molecules^32,33^. When co-cultured with JURKAT cells, *Atg5*-deficient ECs displayed an increased formation of higherJURKAT/EC doublets in a VCAM1-dependent manner indicating that loss of *Atg5* promotes a more proficient interaction of EC with T cells (Extended data Fig. 4f).

**Figure 4:**
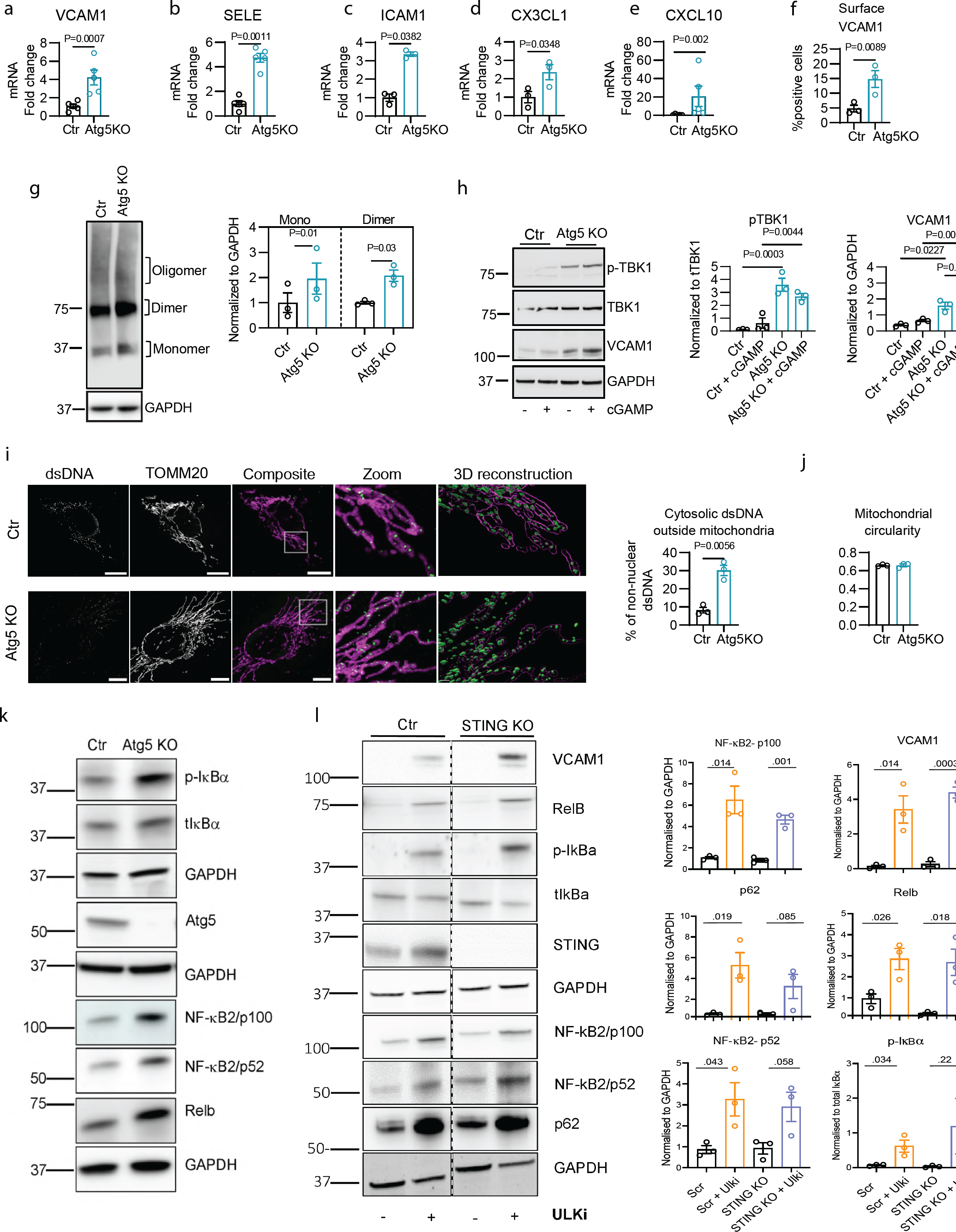
In cultured ECs autophagy curbs inflammation by the CGAS-STING mediated NF-κB pathway through the clearance of cytosolic dsDNA. **a-e,** mRNA expression of *VCAM1* **(a)**, *SELE* **(b)**, *ICAM1* **(c)**, *CX3CL1* **(d)** and CXCL10 **(e)** in and HUVECs after deletion of *Atg5*. **f**, Flow cytometry analysis for surface expression of VCAM1 (% of live cells) in Ctr and Atg5KO HUVECs upon stimulation with IFNγ for 4h. **g,** Representative western blot and quantification of STING monomers, dimers and oligomers in Ctr and Atg5KO HUVECs. **h,** Representative western blot and relative quantification of pTBK1, tTBK1 and VCAM1 in Ctr and Atg5KO HUVECs treated with vehicle or 2.5 μg/mL STING agonist (2’3’-cGAMP) for 24h. **i,** super-resolution Airyscan immunofluorescence images for TOMM20 and dsDNA (green - outside mitochondria, magenta – inside mitochondria) in Ctr and Atg5KO HUVEC. Scale bar represents 10µm. **j,** quantification of mitochondrial circularity per cell in Ctr and Atg5KO HUVECs by super resolution Airyscan immunofluorescence images. Data from HUVECs was generated from at least three independent donors. In all images, scale bars represent 10μm and at least 30 cells (10 per donor) were imaged per condition. **k,** Representative western blot of p-IκBα, total IκBα, Atg5, NF-κB2(p100 and p52), and RelB in Ctr and Atg5KO Huvecs. **i**, Representative western blot and relative quantification of NF-κB2(p100), VCAM1, p62, RelB, NF-κB2(p52), and p-IκBα, in Ctr and STINGKO HUVECs treated or not with ULK1/2 chemical inhibitor (ULK1/2 inhi). All proteins are from the same set of lysates. When proteins were run on the same blot, cropping is denoted by a dashed line) All data show mean ± sem. Statistical difference was determined using a two-sided Student’s t-test **(a-f, i, m)** or one-way Anova with Tukey corrections for multiple comparisons **(g, j)**. Source data are available for **a-m**.

Given the similarity of the *in vivo* and *in vitro* phenotypes, we used cultured autophagy-depleted ECs to gain further insights into the molecular pathways that exacerbate inflammatory signaling. We focused on the STING and NF-κB signaling pathways because STING activation is known to foster antitumor immunity by co-regulating type I IFN and NF-κB activation^30,34,35^, which are enhanced in muTECs of melanoma-bearing Atg5^BECKO^ mice (Fig. 3c,d). Moreover, while autophagy is known to regulate both STING and NF-κB pathways^36^, in ECs, depletion of STING improved the recruitment (however not the adhesion) of T cells in response to TNF-induced inflammation^37^.

In the canonical STING activation pathway, cGAS binds to cytosolic dsDNA and catalyzes the synthesis of cGAMP. Binding of cGAMP to the ER-associated STING triggers its oligomerization, and exit/release from the ER to the ERGIC^38^ where oligomerized STING mediates the activation and phosphorylation of TBK1. This in turn, leads to the transcription of interferon stimulated genes and activation of NF-κB^34^. Signal cessation is orchestrated by STING trafficking from the post-Golgi to lysosomes, where STING is degraded^39^.

We first measured the effects of perturbations of the autophagy/lysosomal degradation pathway on STING trafficking and signaling. In resting conditions, blockade of canonical autophagy either by *Atg5* genetic deletion (Fig. 4g) or pharmacological inhibition of both ULK1/2 kinase activity (Extended data Fig. 4g) led to a significant accumulation of STING both in its monomeric and dimeric/oligomeric forms. Compared to control (Ctr) cells, loss of *Atg5* in ECs increased granular STING staining in the perinuclear regions, a hallmark of STING activation and signaling (Extended data Fig. 4h)^40^. Congruently, in autophagy deficient ECs, STING displayed an increased co-localization with LMAN1 (a marker of the ERGIC compartment) (Extended data Fig. 4h), which was associated with TBK1 phosphorylation (Fig. 4h), suggesting its activation. In line with this observation, addition of cGAMP, the product of the cGAS enzymatic activity, to ECs slightly but not significantly elevated p-TBK1 and VCAM1 in Ctr ECs, whereas it had no additional effects on Atg5KO ECs (Fig. 4h.

We further explored the possibility that autophagy could stimulate the clearance of cGAS activating factors, such as cytosolic dsDNA originating from mitochondria, which is an established endogenous trigger of STING signaling ^41,42^. Autophagy has been shown to inhibit type I IFN in cells undergoing mitochondrial outer membrane permeabilization (MOMP) driven by bona fide apoptotic signals^43–45^. Super-resolution microscopy using TOMM20 to stain the mitochondrial network and an antibody against dsDNA revealed that, compared to Ctr ECs that displayed dsDNA encapsulated within the mitochondrial network (Fig. 4k), dsDNA in Atg5-KO ECs was mostly found in close proximity to the mitochondrial network and partly in the cytosol (Fig. 4i), suggesting its efflux through mitochondrial-derived vesicles^46^. Since we did not observe presence of micronuclei in autophagy-deficient cells (Extended data Fig. 4i), this result further suggests that cystolic dsDNA may be of mitochondrial origin. Although loss of *Atg5* in ECs compared to Ctr ECs did not induce evident changes in key mitochondrial morphometric parameters (Fig. 4j) or alterations in the amount of mitochondrial proteins (Extended data Fig. 4d) as a measurement of mitochondrial clearance; we cannot exclude that a minority of mitochondria undergo MOMP-mediated mitochondrial dsDNA release^43,47^, or other forms of mitochondrial stress, when autophagy is dysfunctional. Also, given that STING harbors an LC3-intercating regions (LIR) and can be targeted for autophagic degradation^48,49^, we cannot exclude that this mechanism contributes to the STING signal observed in *Atg5*-depleted ECs. Notwithstanding, these results suggest that loss of autophagy in ECs mediates cGAS-STING and NF-κB activation, resulting in the expression of a repertoire of inflammatory/immunomodulatory target genes. Congruently, *Atg5* ablation in ECs along with the upregulation of VCAM1, led to the phosphorylation of the inhibitory IκBα protein (RelA) of the canonical NF-κB pathway, and increased the accumulation of mediators of the alternative NF-κB signaling^50^; the precursor protein p100 and its degradation product p52, and RelB, suggesting the activation of both canonical and alternative NF-κB pathways (Fig. 4k)^51^. This is in line with the upregulation of multiple regulators of the NF-κB pathways, including *RELB* upon deletion of *Atg5* in murine TEC (Fig. 3c).

We then asked whether cGAS-STING was required for NF-κB activation in autophagy-depleted ECs. To this end we CRISPR-Cas9 deleted STING in Ctr and in autophagy inhibited cells. However, the effects of the concomitant and chronic suppression of Atg5 and STING varied across human EC donors making these results unreliable. We then tested the effects of ULK1/2 inhibition in STING-depleted cells. In analogy with *Atg5* depletion, ULK1/2 inhibition elicited the upregulation of VCAM1, IκBα-phosphorylation and the accumulation of p100, RelB and p52, and increased the presence of autophagic receptor p62, which is degraded by autophagy (Fig. 4l). The effects of ULK1/2 inhibition on these effectors were still retained in STINGKO ECs (Fig. 4l), suggesting that STING is dispensable for the activation of NF-κB pathways in autophagy-depleted ECs.

In summary, endothelial autophagy blockade enhances the immunosupportive/inflammatory TEC phenotype by the activation of both STING signaling and STING-independent NF-κB pathways.

### LOSS OF Atg5 IN TUMOR ENDOTHELIAL CELLS SUSTAINS NF-kB-DRIVEN INFLAMMATION INDEPENDENT OF STING

Previous studies, using conditional whole-body STING KO mice or STING activation by intratumoral injection of STING agonists, proposed that vascular STING expression and Type I IFNs promote antitumor immunity^5,35^. Considering that species-specific mechanisms control the strength and specificity of STING downstream signaling responses^52^, we then sought to delineate the contribution of STING signaling to melanoma growth control in WT and Agt5^BECKO^ mice.

To this end, we crossed EC-specific *Sting* KO mice^53^ with Atg5^BECKO^ mice to generate EC specific conditional *Atg5/Sting* double knockout (ATG5/STING^BECDKO^) mice (Extended data Fig. 5a,b).

**Figure 5:**
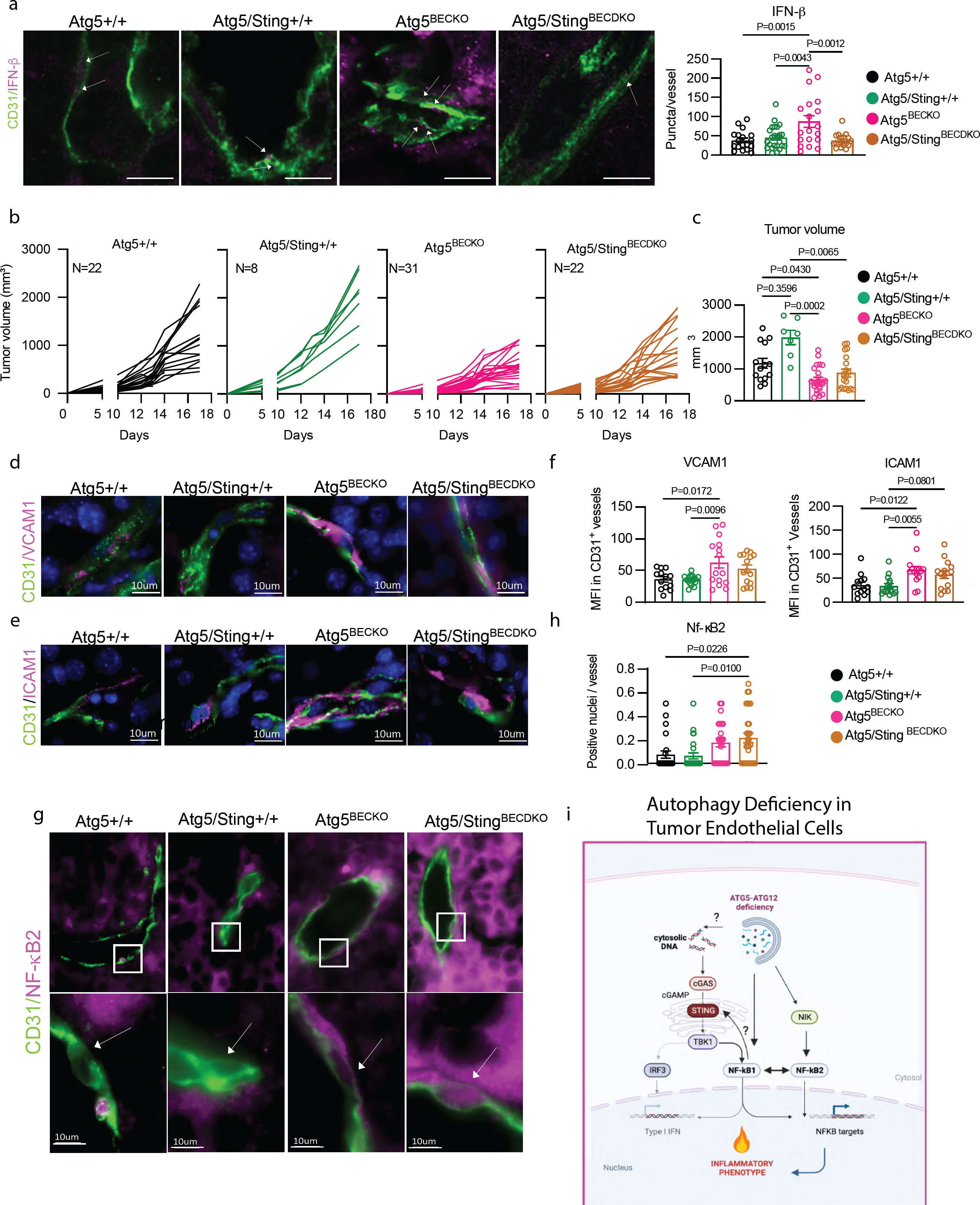
Dual genetic deletion of *Sting* and *Atg5* in TECs promotes the activation of the alternative NF-κB pathway. **a,** Representative images and quantification (puncta per vessel) of immunofluorescence staining for IFNβ in the tumor sections of subcutaneous B16-F10 tumors from Atg5+/+, Atg5/Sting+/+, Atg5^BECKO^ and Atg5/Sting double KO (Atg5/Sting^BECDKO^) mice. Scale bar represents 10µm**. b-c,** Individual tumor volume **(b)** and end-point tumor volume **(c)** of Atg5+/+, Atg5/Sting+/+, Atg5^BECKO^ and Atg5/Sting^BECDKO^ mice subcutanesouly injected with B16-F10 melanoma cells. Figures are pooled from two independent experiments **(b-c)**. **d-f,** representative images and quantification (MFI) of immunofluorescence staining for CD31 **(d,e,)**, VCAM1 (**d)**, ICAM1 **(e)** in the tumor sections of subcutaneous B16-F10 tumors from Atg5+/+, Atg5/Sting+/+, Atg5^BECKO^ and Atg5/Sting^BECDKO^ mice. **g,h,** representative images and quantification (Positive nuclei) of immunofluorescence staining for CD31 and NF-кB2 in the tumor sections of subcutaneous B16-F10 tumors from Atg5+/+, Atg5/Sting+/+, Atg5^BECKO^ and Atg5/Sting^BECDKO^ mice. Scale bar represents 10µm. **i**, schematic of proposed model for inflammatory phenotype downstream of autophagy inhibition. For immunofluorescence staining, at least 3 mice per group were used for the quantification. All data show mean ± sem. Statistical differences were determined by one-way Anova with Tukey corrections for multiple comparisons. Source data available for **a-g**.

We then injected B16-F10 melanoma in Atg5^+/+^ mice, Atg5/STING^+/+^, Atg5^BECKO^ and Atg5/STING^BECDKO^ mice, monitored tumor growth and inspected the TEC phenotype by IHC for the presence of proinflammatory/immunomodulatory factors.

The granular expression of IFNß, a major downstream target of STING signaling, was elevated in TECs from melanoma-bearing Atg5^BECKO^ mice, whereas its expression was blunted in the tumor vessels of Atg5/STING^BECDKO^ mice and comparable to that of the WT TECs (Fig. 5a), thus functionally validating the TEC phenotype of the DKO mice.

Melanoma growth (Fig. 5b-c) was significantly reduced in both Atg5^BECKO^ and Atg5/STING^BECDKO^ mice and did not differ in the recruitment of intratumoral CD3+ T cells (Extended data Fig. 5c). Moreover, double staining of CD31 and VCAM1 or ICAM1, showed that TECs from both Atg5^BECKO^ and Atg5/STING^BECDKO^ B16F10 melanoma-bearing mice displayed elevated levels of VCAM1 and ICAM1 adhesion molecules compared to TECs from their respective WT melanomas/controls (Fig. 5d-f). Since VCAM1 and ICAM1 are shared downstream targets of the canonical and alternative NIK-mediated RelB/p52 pathways^54^ (Fig. 3c, Fig. 4k,l), we analyzed the levels of NIK and of the nuclear localized/active p52 transcription factor in TECs from Atg5^+/+^, Atg5/STING^+/+^, Atg5^BECKO^ and Atg5/STING^BECDKO^ tumor-bearing mice. Compared to their respective WT counterparts, TECs from both Atg5^BECKO^ and Atg5/STING^BECDKO^ displayed increased levels of NIK (Extended data Fig. 5d). Interestingly, concomitant EC-specific deletion of both *Atg5* and *Sting* (i.e., in the Atg5/STING^BECDKO^) (Fig. 5g,h) exhibited a trend toward further increase in CD31+ vessels displaying p52 positive nuclei when compared to single *Atg5* deletion (i.e. Atg5^BECKO^ mice), suggesting that the alternative NF-κB pathway was still heightened in the absence of STING, in sync with the *in vitro* data (Fig. 5i).

Together, these data posit that STING signaling in autophagy-deficient TEC contributes to the exacerbating NF-κB-dependent immunosupportive TEC phenotype, but is not essential.

### TUMOR ENDOTHELIAL AUTOPHAGY LIMITS MELANOMA RESPONSES TO ANTI-PD1 THERAPY

Given the immune-stimulating role of autophagy blockade concomitant with enhanced CD8 T cell infiltration and activity, we then inquired whether impairing autophagy in TECs can boost immunotherapy in the form of immune-checkpoint blockade (ICB) (Fig 6a).

**Figure 6:**
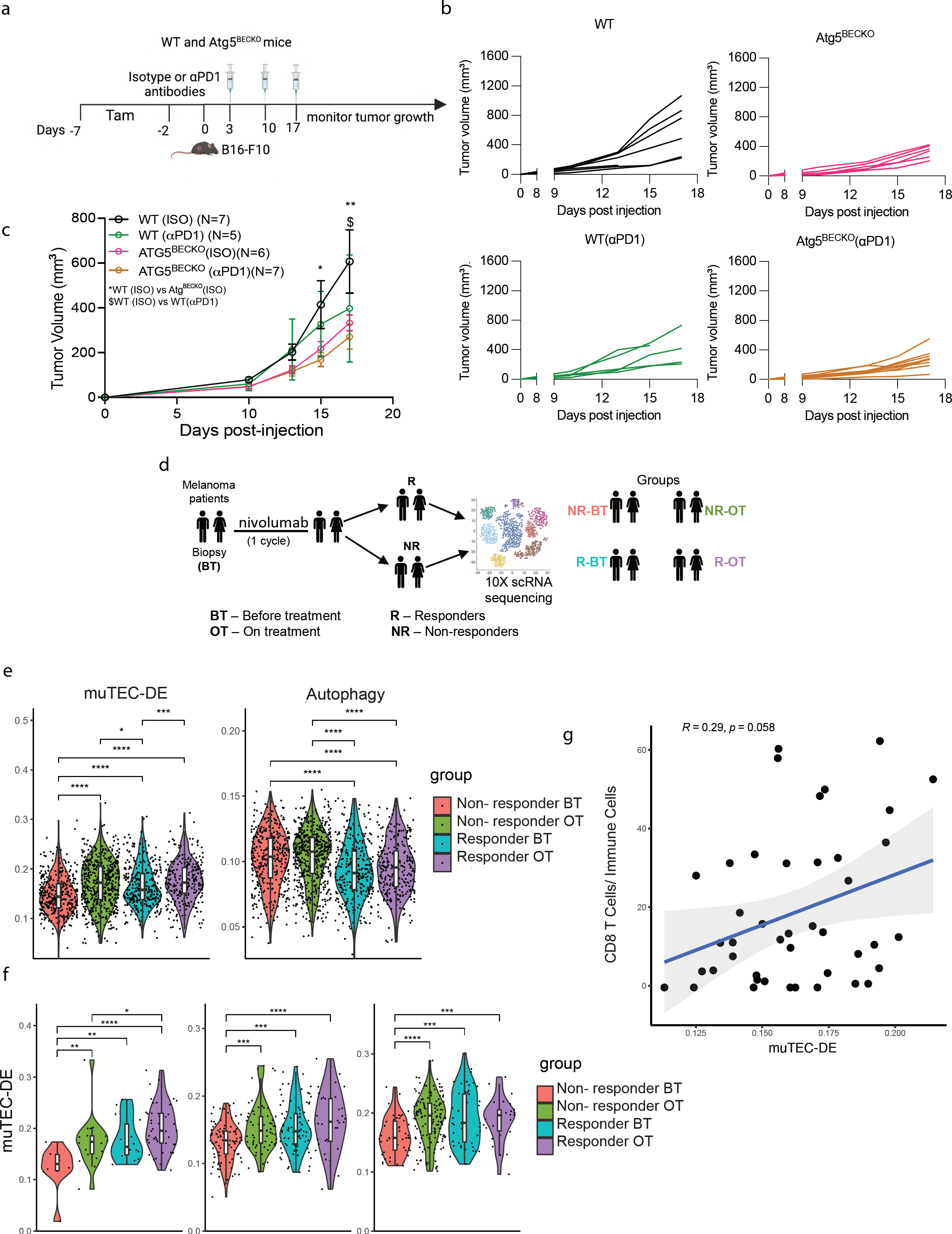
TEC-autophagy negatively correlates to the response to anti-PD1 therapy in melanoma. **a,** Schematic representation for treating subcutaneous B16-F10 tumor bearing WT and Atg5^BECKO^ mice with αPD1 therapy. **b-d,** Individual tumor volume **(b)**, grouped tumor volume **(c)** of B16-F10 subcutaneous tumor bearing WT and Atg5^BECKO^ mice injected with isotype (ISO) or αPD1 antibody. Data shown in **b,c** are from 1 representative experiment **d,** Study design for single-cell transcriptome analysis of huTEC from treatment naïve stage III/IV melanoma patients receiving anti-PD-1 based therapy monotherapy (nivolumab). **e-g**, scRNA-seq data analysis from huTECs in melanoma. **e**, AUCell enrichment score of muTEC-DE (extended data table 1)and autophagy (Reactome database, R-HSA-1632852: Macroautophagy) genesets in all ECs. **f**, AUCell enrichment score of mTEC DE geneset in 3 EC subtypes. **g** correlation between AUCell score of mTEC DE geneset and percentage of CD8 T cells in each sample. Median-quantile-min/max + population distribution is shown in violin + boxplot (f, g). Wilcoxon test, ∗p < 0.05; ∗∗p < 0.01; ∗∗∗p < 0.001; ∗∗∗∗p < 0.0001. All data represent mean ± sem (**c)**. Statistical differences were determined by two-way Anova with multiple comparisons *$ P < 0.05, ** P < 0.01. Source data available for **a-c**.

While anti-PD1 monotherapy significantly reduced B16-F10 tumor growth in WT mice (Fig. 6b,c), the effect of immunotherapy in B16F10 bearing Atg5^BECKO^ mice, was incremental but not significant (Fig. 6b,c). After cessation of anti-PD1 therapy tumor burden in ICB-treated melanoma-bearing WT mice rapidly resumed (Extended data Fig. 6a), whereas in Atg5^BECKO^ treated with anti-PD1, or their isotype Atg5^BECKO^ controls, tumor burden was more efficiently contained (Extended data Fig. 6a). These results suggest that blockade of autophagy in TEC of melanoma-bearing mice, by favoring recruitment and activity of TILs (Fig. 2d-g), does not further enhance the efficacy of anti-PD1 ICB therapy, but it may further sustain its effects.

Our previous analysis indicated that particularly the huTEC interferon subtype of treatment-naïve cancer patients exhibited a divergent association between the expression of muTEC-DE^High^ and autophagy^Low^ gene signatures (Fig. 3f). To further portray the significance of these TEC-specific gene signatures and their association with clinical responses of melanoma patients to ICBs, we then interrogated the single-cell transcriptome of huTECs using scRNAseq data from a unique prospective longitudinal study, including treatment naïve stage III/IV melanoma patients receiving anti-PD1 based therapy. Tumor biopsies were collected before (BT) treatment and right after the first cycle of treatment (ON treatment) and processed for scRNAseq analysis.

Data consisted of TECs from 22 samples, N=12 responders (R) and N=10 non-responders (NR) (Fig.6d). Overall, we annotated 1541 ECs using established methods for unbiased EC identification^55^. Of note, the melanoma EC data set included several EC cell types which were absent from the other cancer types (Fig. 3). Melanocyte-like cells are unique to melanoma, and may exist here due to vasculomimicry. In addition, pericytes were included in this dataset. (Extended data Fig. 6b-c). Compared to non-responders (NR), huTEC from responders (R) showed a significant enrichment of the core inflamed gene sets (muTEC-DE) before (BT) treatment (Fig. 6e), while showing a lower enrichment in the autophagy gene core signature. Interestingly, non-responders (NR) also showed an enrichement in the muTEC-DE gene signature after one cycle of treatment (Fig.4e). This might suggest that a TEC inflammatory phenotype is associated with an initial (yet not long-lasting and therapeutically inefficient) change in the TME driven as a first response to anti-PD1 immunotherapy. However, in the same set of patients the higher autophagy score was unchanged (Fig.4e), suggesting that only the muTEC^High^/autophagy^Low^ status is associated with clinical benefits.

We then interrogated whether muTEC-DE gene signature differed between responders (R) and non-responders (NR) exclusively in huTECs subtypes. To do so we clusters all huTECs and obtained 11 subtypes reflecting huTECs heterogeneity observed in primary pan-cancer data (Extended data Fig. 6a (UMAP), b (heatmap)). Compared to non-responders (NR), melanoma patients responding to anti-PD1 harbored a significant enriched muTEC-DE gene signature in the Interferon, Tip/Stalk and Venous EC clusters, a trend that was stably maintained at the ON treatment time point in the Interferon subtype (Fig. 6h; Extended data Fig. 6c). Furthermore, using these scRNAseq data sets, we found a positive correlation between the frequency of the muTEC-DE signature in all TECs and the intratumoral infiltration of CD8+ T cells (Fig. 6g).

Together, these data suggest that the muTEC-DE ^High^/autophagy^Low^ phenotype is clinically associated with a favorable response to anti-PD1 therapy.

### SPATIAL PROXIMITY BETWEEN INFLAMED VESSELS AND CD8+ T CELLS CORRELATES TO RESPONSE TO ANTI-PD1 THERAPY

We then sought to portray the spatial relationship between inflamed huTECs as defined by their enhanced protein expression of VCAM1 or double positive VCAM1 and STING, and CD8+ T cells in melanoma samples. We performed multiplex immunohistochemistry with the multiple iterative labelling by antibody neodeposition (MILAN) technology^56,57^ (Extended data Fig. 7a-c) using a set of real-life biopsies from melanoma patients undergoing anti-PD1 monotherapy (extended data table 2). The melanoma biopsy cohorts consisted of 6 responders (R) and 6 non-responders (NR). CD31/AQP1 double positive (identifying huTECs), CD8+ PD1-GrzB-T cells (identifying naïve CD8+ T cells) and CD8+ PD1+ GrzB+ (identifying activated/effector CD8+ T cells)^58^ were evaluated in three distinct spatial compartments of the biopsy, namely the “tumoral bulk”, the tumor-stroma interface, and the non-tumoral areas. There was no difference in EC number between non-responders (NR) and responders (R) in the the tumor bulk and at the tumor-stroma interface (Fig. 7a,b). We then investigated the proximity among these cell types in these spatial compartments by neighbourhood analysis. In contrast to the non responders (NR), responders (R) showed a significant enrichment of VCAM1^+^ and VCAM1^+^STING^+^ TECs in the tumor bulk and at the tumor-stroma interface, respectively (Fig. 7c,d). Further neighbourhood analysis showed that responders (R) had a significantly higher number of VCAM1^+^STING^+^ TECs in vicinity to both of CD8+ PD1-GrZB- and to CD8+ PD1+GrZB+ T cells (Fig. 7e,f). This observation was true for the tumor-stroma interface and the tumor bulk. Particularly within the tumor bulk, even if in lower number, VCAM1^+^ vessels were still significantly closer to both subsets of CD8+ T cells, suggesting that an inflamed vessel phenotype maintain a more proficient crosstalk with infiltrated TILs (Fig. 7g,h), even in the absence of STING. We also used LC3 antibody in the MILAN analysis, but due to limited resolution of the intracellular granular vs diffuse pattern of LC3 staining in TECs we could not reliably assess their associated autophagy status.

**Figure 7:**
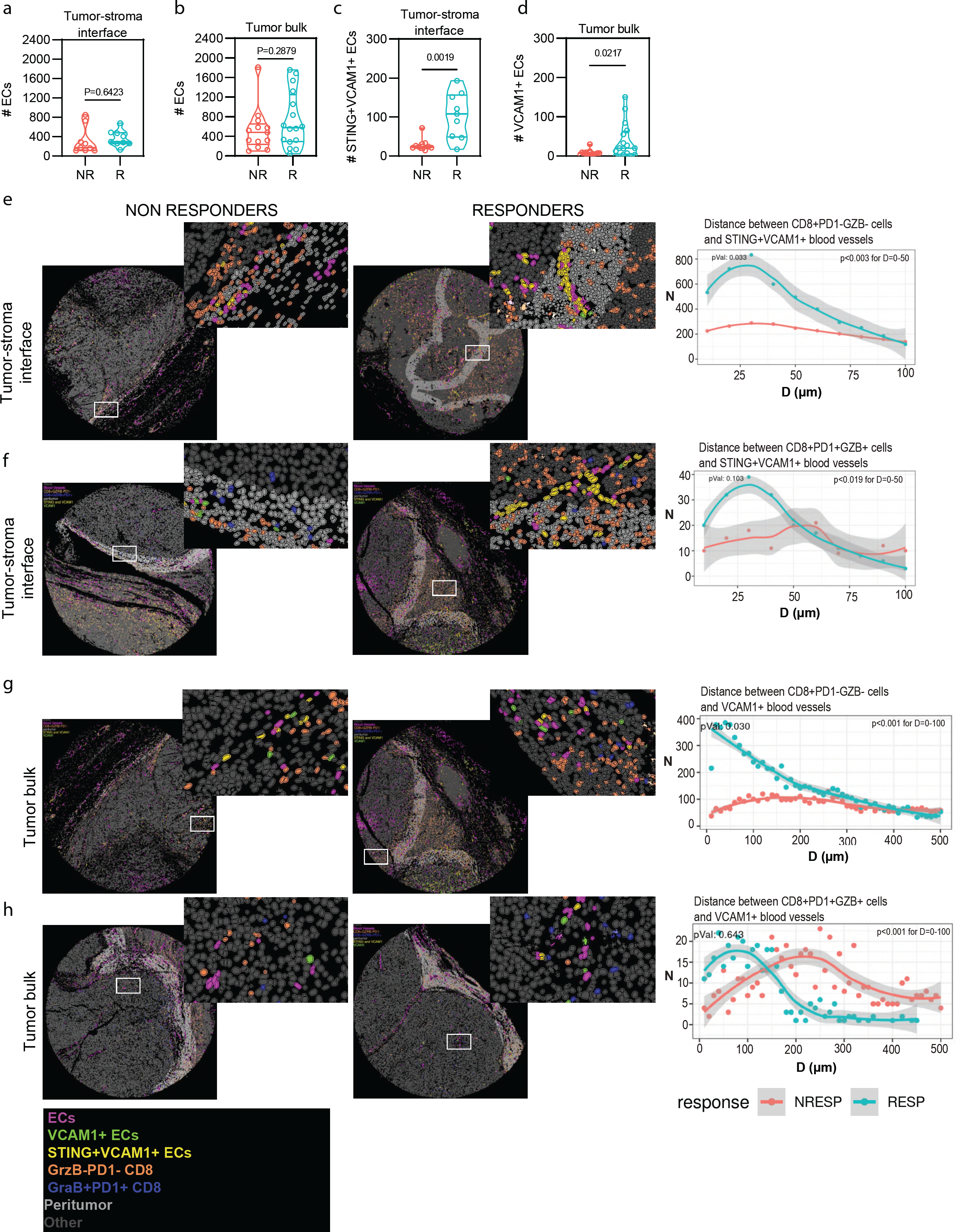
Responders to anti-PD1 therapy house inflammed TECS with immune cell attracting properties. **a-d,** Violin plots comparing non-responders (NR) and responders (R) in terms of **(a)** total number of ECs at the tumor-stroma interface **(b)** total number of ECs in the tumor bulk, **(c)** number of VCAM1^+^STING^+^ ECs in the tumor-stroma interface **(d)** number of VCAM1+ ECs in the tumor bulk. **e-h,** representative digital reconstruction of the tissue based on multiplex staining and cell clustering illustrating EC subtypesand CD8^+^ T cells subtypes. The results of the neighborhood analysis between ECs and CD8^+^ T cells subtypes are visualized by the line graph on the right. Neighborhood between ECs and CD8+ T cells subtypes is expressed as number of cells (y axis) located at a certain distance (x axis) from the ECs. All data represent mean ± sem. *P < 0.05, **P < 0.01, ***P < 0.01, ****P < 0.0001 using a two-sided Student’s t-test **(a-f)**. In neighborhood analysis of CD8^+^ T cells with ECs **(e-h)**, two-sided Student’s t-test was used at the particular distance. Source data are available for **a-f**.

Together these results show that the spatial proximity between inflamed vessels with infiltrating CD8+ T cells is a hallmark of the TME of melanoma patients responding to anti-PD1 therapy.

## DISCUSSION

Despite the emerging evidence indicating that tumor blood vessels are not just passive bystander regulators of immune cell trafficking, but play an active role in antagonizing antitumor immunity^8,27,59^, the mechanisms controlling the immunosuppressive phenotypes of tumor ECs remain largely unexplored. Gaining mechanistic knowledge of the processes that remodel the tolerogenic/anergic phenotype of TEC into a more proinflammatory/immunostimulatory status is thus key to improve immunosurveillance and immunotherapy responses. The primary focus of this study was to investigate the molecular mechanisms endorsing the tolerogenic status of tumor endothelial cells (TECs), and to functionally assess the consequences of perturbing these mechanisms on melanoma antitumor immunity.

The data presented in this study advance the concept that autophagy is a crucial vascular-immune checkpoint that endorses TEC-mediated immune cell barrier and immunoevasion.

First, through the EC-specific ablation of assorted autophagy genes (*Atg5, Atg12* and *Atg9)* involved in the early phases of autophagosome formation and expansion, we show that impairing autophagy reduces melanoma growth by promoting expansion of effector CD4+ and CD8+ T cells, thereby reducing the immunosuppressive TME. Second, we unveil that the inhibition of TEC-autophagy potentiates the proinflammatory/proadhesive and immunomodulatory functions of TECs, by exacerbating the canonical and alternative NF-κB pathways, and by promoting STING signaling. Mechanistic and functional studies further highlight that NF-κB activation is the major driver of the pro-inflammatory TEC phenotype induced by loss of autophagy. Third, by interrogating scRNAseq EC atlases of melanoma patients undergoing one cycle of anti-PD1 immunotherapy, we show that TEC hallmarked by an Inflammatory^High^/autophagy^Low^ phenotype correlates with clinical responses to anti-PD1. In line with this, by using multiplex immunohistochemistry, we unravel that melanoma patients responding to anti-PD1 therapy present inflamed vessels in close neighborood with naïve and activated CD8+ T cells.

Autophagy has a well established role in the suppression of homeostatic and pathological inflammation^60^. Due to its innate ability to clear intracellular (principally mitochondria-derived danger signals) and extracellular (pathogenic) cargos^61^. It also regulates differentiation and metabolic programs in a highly context and cell or tissue-specific manner^61^.

In cancer, recent studies have shown that conditional whole-body or hepatocyte-specific knockout of *Atg5* or *Atg7* improve antitumor immunity by stimulating IFN-γ-mediated increase in MHC class I expression, and antigen presentation of murine tumors with high mutational burden, through STING-mediated immunity^62,63^. While these studies implicated the existence of multiple signaling circuitries controlled by host autophagy that regulate antitumor immunity, it remained unclear whether autophagy, specifically in the tumor endothelium, would be a major regulator of these immunosuppressive responses and if it could affect the efficacy of immunotherapy.

Our transcriptomic profiling of TECs isolated from WT and Atg5^BECKO^ melanoma-bearing mice, leveraged an inflammatory signature -dubbed muTEC-DE-which we then used to probe publicly available TEC taxonomies from different primary human tumors types. Using this in-house generated signature we found that across previously annotated TEC subtypes^12,55^, the enrichment of the muTEC-DE signature was associated with the lowest expression levels of autophagy genes, particularly in the Interferon EC subtype. This divergent association, which conforms to our functional data, raises the question of how particularly this subtype of TEC maintains a low transcriptional expression of autophagy genes. Recent scRNAseq studies obtained from different tumor types have highlighted that metabolic plasticity is an emerging trait of the phenotypic adaptation of TECs to their specific TME^64^. Supporting a link between metabolism/epigenetic and IFN response, loss of the glycolytic enzyme PKM2 in ECs by altering the TCA cycle, have been shown to promote DNA hypomethylation, de-repression of endogenous retroviral elements, with the resulting activation of antiviral innate immune signalling^65^. Hence, it is tempting to suggest that autophagy gene expression may be co-regulated as a result of different metabolic/epigenetic circuits adopted by a specific EC phenotype. This is an outstanding question that warrants further investigation.

Consistent with the results in murine melanoma and other human tumors, in treatment-naïve melanoma patients the TEC subtype displaying the muTEC^High^/autophagy^Low^ phenotype was associated to clinical responses and to a higher numbers of tumor infiltrating CD8+ T cells, further suggesting the functional association between TEC-autophagy and suppression of immune-related responses. Furthermore, compared to non-responders, patients responding to anti-PD1 monotherapy had an increased VCAM1+STING+ vessel counts found in close vicinity to CD8 + T cells and the fraction of effector PD1+/GrzB+ CD8+ T cells, particularly at the tumor-stroma interface. A similar positive correlation was found with VCAM1+ vessel counts within the tumor bulk. These findings are congruent with the observation that elevated STING expression in tumor vessels from colorectal cancer patient biopsies correlated with the intratumoral CD8+ T cell infiltration^5^. A recent study across different cancer types including melanoma, indicated that an elevated frequency of CD8+ T cells within the stroma and invasive margin compartments had a better outcome than those in intra-tumor compartments^66,67^. However, more studies are needed to assess how positioning of the inflamed vessels within the tumor parenchyma models their immunomodulatory phenotypes (tolerogenic versus immunostimulatory) and whether the IFN-EC subtype in responders is accrued at the tumor-stroma interface to facilitate the infiltration of CD8 +T cells.

But which signals are critical to convey the proinflammatory and antitumor effects of autophagy ablation in TECs? Recent studies have largely focused on the crosstalk between autophagy and STING pathways and their effects on cancer cell immunogenicity and antitumor immunity in response to therapy^68,69^. However, emerging evidence indicate that the homeostatic and pathological functions of autophagy are shaped by the cellular and tissue/microenvironment context^19,26^ and respond to different inflammatory cues. Previous studies indicated that within the tumor microenvironment, TECs are early producer of Type I IFN, following intratumoral injection of cGAMP, and contribute via STING to the initiation of spontaneous and ICB induced antitumor immunity^35^. Exogenous, agonist-mediated STING activation induces vascular reprogramming, likely caused by the reciprocal beneficial effects of tumor-infiltrating CD8+ T cells on the tumor vasculature^5,70^. However, monotherapy with STING agonist, which sustains Type I IFN signaling, may also elicit negative feedback mechanisms of resistance^71^, requiring the concomitant combination of anti-angiogenic (anti-VEGFR2 blockers) and ICBs (anti-PD1 or anti-CTLA4) to be effective^5^.

While these studies highlight the relevant function of the tumor vasculature in antitumor immunity, they did not address the underpinning mechanisms of STING modulation specifically in ECs, nor the role of autophagy as potential mitigation signal. Here, we report that upon ablation of *Sting* in *Atg5*-deficient TECs (EC-specific *Atg5/Sting* double knockout) the stimulation of NF-κB pathways maintains the proadhesive/inflammatory TEC phenotype. This is likely the reason why tumor burden in STING/Atg5^BECDKO^ mice is still reduced, to an extent that is similar to what observed in *Atg5*-deficient TECs. While we cannot exclude that concomitant genetic loss of *Atg5* and *Sting* in muTECs have additional cell-non autonomous effects, which need to be further addressed, our *in vitro* and *in vivo* data, indicate that STING is not required for the activation of NF-kB and is dispensable for the major inflammatory effects of autophagy inhibition in TECs, since in the absence of STING, NF-κB pathways remain active. Interestingly, while STING is known to mediate the activation of NF-κB by recruiting TBK1^34^, recent data posited that NF-κB activation, by inhibiting microtubule-mediated STING transport to the lysosome, promotes STING signaling in response to a variety of signals^72^. These studies unravel the existence of a complex positive feedforward loop between these immunostimulatory pathways, which is likely regulated in a cell dependent manner. The exact mechanisms linking the possible co-regulated activation of STING and NF-κB proinflammatory pathways in response to endothelial autophagy inhibition, still need to be further explored. The finding that the expression of *RELB* is upregulated in *Atg5*-deficient TECs isolated from melanoma-bearing mice, suggest a transcriptional mechanism. However defects in the turnover of adaptors^73–76^ or elements of canonical and noncanonical NF-κB pathways, caused by the accumulation of undegraded p62 or other autophagic adapter proteins, could support NF-κB proinflammatory signals under conditions of TEC-autophagy suppression^54^. Notwithstanding, our results show that autophagy is able to repress two major immunomodulatory pathways, NF-κB and STING, in tumor vessels thus endorsing their immunosuppressive phenotype, with deleterious consequences for antitumor immunity and efficacy of ICBs.

Considering the broader and translational relevance of our findings, this study supports the emerging concept that strategies aimed to suppress the anergic status of tumor endothelial cells are decisive to overcome immunoevasion and to improve efficacy of various immunotherapy approaches^77^. As found in other tumors^5^, in preclinical models of melanoma intratumoral STING agonist (i.e., ADU S-100) enhances tumor infiltrating T cells and reduced melanoma growth^35,78^, depending on the host, not melanoma cell expression, of STING. The results of our study advocate that TEC-autophagy blockade, by the concomitant and independent activation of STING and canonical and noncanonical NF-κB pathways, offers a more efficient strategy to overcome immunoevasion in melanoma. Therefore, it will be important to scrutinize the still unappreciated effects of clinically used autophagy inhibitors^79^ on the tumor vasculature. Previous studies from our laboratory showed that the clinically used lysosomotropic drug chloroquine (CQ), exerted antimetastatic effects in melanoma by primarily normalizing the tumor vasculature through enhanced Notch signaling^22^ but the effects of CQ on NF-κB signaling in TECs were not explored. However, systemic administration of CQ could have additional and possible indirect effects on T cells or other stromal cells, making it the clinical utilization of this lysosomotropic drug as a vascular autophagy inhibitor problematic.

In summary, this study advances our knowledge on the role of autophagy as a persuasive blood vessel-intrinsic anti-inflammatory and immunosuppressive mechanism, restricting antitumor immunity in melanoma. It also provides a theoretical foundation for identifying appropriate combination therapies utilizing vasculature-homing tools^80^ to target autophagy inhibitors or NF-κB modulators^81^ to the tumor vasculature.

## METHODS

### Cell Culture

#### Primary cells

HUVECs from different donors were purchased from PromoCell. Cells were maintained in supplemented Endothelial Cell Growth Medium 2 kit (PromoCell #C-22111) at 37 °C under 5% CO2.

#### Murine Melanoma cell lines

B16-F10 and HcMEL12-mCherry cells were maintained in RPMI-1640 (Sigma #R8758) medium supplemented with 10% FBS, 1mM sodium pyruvate and 10mM HEPES at 37 °C under 5% CO2. Yummer 1.7 were maintained in DMEM/F12 medium (#DF-041-B) supplemented with 10% FBS and 1X non-essential amino acids (#TMS-001-C) and pen/strep.

#### Inhibitors, agonists and cytokine

ULK1 and ULK2 Kinase inhibitor MRT68921 (0.5 μM; Sigma #SML1644-5MG), STING agonist 2’3’-cGAMP (2.5µg/mL; Invivogen# tlrl-nacga23), TNFα (5 ng/m; R&D systems #210-TA-020) and IFNγ (25 IU/mL; Sigma #SRP3058-100UG) were used as recommended by the supplier.

### CRISPR-Cas9 gene knockout by nucleofection

Genes of interest were deleted by nucleofecting Ribonucleoprotein complexes (RNPs) consisting pooled sgRNAs conjugated with Cas9. Nucleofection was performed using the P5 Primary Cell 4D-Nucleofector kit (Lonza # V4XP-5024). RNPs were prepared by co-incubating 500 pmoles of sgRNA pools (for Atg5, cGAS and STING) with 100 pmoles of SpCas9 2NLS (Synthego, USA) and introducing the RNPs into 700,000 HUVECs using CA-167 programme in Lonza 4D nucleofector. Cells were allowed to recover for 48h before manipulating them for further experiments.

### Mice

All experimental animal procedures were approved by the Institutional Animal Care and Research Advisory Committee of the KU Leuven (PC57BL6). Mice with blood endothelial cell (BEC) specific deletion of *Atg5* were obtained by crossing Pdgfb-Cre^ERT2^ Rosa26^tdTomato/tdTomato^ mice^82^ with previously generated *Atg5*^fl/fl^ mice^83^. Mice expressing Cre (Pdgfb-Cre+^ERT2^; Atg5^fl/fl^) were referred to as Atg5^BECKO^ or their Cre-negative littermates were referred to as WT. Mice with pan endothelial cell (EC) specific deletion of Atg12 or Atg9a were obtained by crossing mice VeCadh-cre^ERT2^ ^84^ with *Atg12*^fl/fl8569^ or *Atg9a^fl/fl^* mice^86^. Mice expressing Cre (chd5-Cre+^ERT2^; *Atg9a*^fl/fl^ or *Atg12*^fl/fl^) were referred to as Atg9a^ECKO^ or Atg12^ECKO^ and their Cre-negative littermates were referred to as WT. *Sting*^fl/fl53^ mice were a gift from Dr. Hamida Hamad and Dr. Jonathan Maelfait labs and were crossed with Atg5^BECKO^ mice to obtain conditional Atg5/STING double knockout (DKO) (Pdgfb-Cre^ERT2^; Atg5^fl/fl^STING^fl/fl^) Atg5/Sting^BECDKO^ mice. The Cre-negative littermates were mice were maintained as controls. Both female and male mice from 7 to 14 weeks of age were used for experiments. Inducible deletion was obtained by peritoneally injecting them with tamoxifen (Sigma # T5648-5G) dissolved in cornoil (20mg/mL; Thermo # 405435000). Tamoxifen was injected once a day for 5 days. For tumor cells, subcutaneous injection of 250.000 tumor cells (B16-F10, HCMEL12-mCherry, YUMMER1.7 at 50-60% confluency) suspended in 100µL of PBS was performed under isoflurane.

### CD8^+^ T cell depletion

CD8+ T cells were depleted by intraperitoneal administration of 100 μg of αCD8 antibody (Clone YTS-169) in 100µL PBS three days before B16-F10 tumor cell injections. αCD8 injections repeated on the day of tumor cell injection and on day 5, 10, 14 and 21 after tumor cell injection. 14 mice received anti-CD8 therapy (8 WT and 6 KO mice) and 14 mice received PBS as control (6 WT and 8 KO mice).

### Anti-PD1 treatment

αPD1 antibody (100 μg in 100µL; clone – RMP1-14) were administered intrapertonially three days after B16-F10 tumor cell injections. Treatment was repeated on the day 10 and 17. 21 mice received anti-PD1 therapy (9 WT and 12 Atg5^BECKO^mice) and 19 mice received αbetaGAL rat IgG2a isotype control antiobdy (Clone – GL117) as control (9 WT and 10 Atg5^BECKO^ mice). Both αCD8 and αPD1 antibodies were a kind gift from Dr. Louis Boon (Polpharma Biologics).

### Flow cytometry

#### Murine Tumor Endothelial Cells (muTECs)

For sorting muTECs, subcutaneous tumours were post-morten excised, digested and sorted using method published^87^. For Surface and intracellular staining of muTECs, cells after magnetic encirhment of CD31+ were used. Cell surface immunostaining was done with following antibodies: CD31 (1:400 dilution, BV421; BD #562939), CD45 (1:200 dilution, APC-Cy7; BD #561037), VCAM1 (1:200 dilution, FITC; Biolegend #105705), ICAM1 (1:200 dilution, PE; BD #568539), MHCI (1:100 dilution, APC; Biolegend #107613) and MHCII (1:100 dilution, FITC; Biolegend #107605). eBioscience™ Fixable Viability Dye eFluor™ 780 (1:1000 dilution,Thermo # 65-0865-14) was used for discriminating dead cells. For intracellular staining of IFNβ, cells were fixed and permebilized following the protocol described in eBioscience™ Foxp3/Transcription Factor Staining Buffer Set. Cells were incubated overnight at 4°C with anti IFNβ antibody (1:50 dilution; Thermo #PA5-20390) conjugated to PE in-house by PE/R-Phycoerythrin Conjugation Kit - Lightning-Link (abcam #ab102918). Data was acquired in BD symphony A5 flowcytometer.

#### Immunophenotyping

CD45+ cells obtained during muTEC enrichment were used for immunophenotyping. After selection, 2*10^6 cells were resuspended in DPBS and incubated with eBioscience™ Fixable Viability Dye eFluor™ 780 (Thermo #65-0865-14) for dead cell discrimination. Nonspecific binding of antibodies to cell Fc receptors was blocked using 10 μl FcR blocking reagent (BD Purified Rat Anti-Mouse CD16/CD32 553142) per 10^7^ cells. Cell surface immunostaining was done with the following antibodies: CD45 (1:400 dilution, BUV805; BD #741957), CD3 (1:100 dilution,PE-Cy7; BD#560591), CD4 (1:100 dilution, BUV395; BD #565974), CD8a (1:200 dilution, BUV496; BD #750024), TIM3 (1:100 dilution, BB515; BD #567810), PD1 (1:100 dilution, BV421; BD #562584). After staining with surface proteins, cells were fixed and permeabilized following protocol described in eBioscience™ Foxp3/Transcription Factor Staining Buffer Set. After permeabilization, cells were immunostained with the following antibodies: TCF-7/TCF-1 (1:50, AF647; BD #566693), Granzyme B (1:75, PE; Miltenyi Biotec #130-116-486) and Ki67 (1:50, PerCP Cy5.5; BioLegend #652425). Data was acquired in BD Symphony A5 or SONY ID7000 spectral flowcytometer.

#### HUVECs

Cells were stimulated with IFNγ for 4h and the trypizinized cells were immunostained with VCAM1 antibody (1:100 dilution, BV421; BioLegend #305815). eBioscience™ Fixable Viability Dye eFluor™ 780 was used for discriminating dead cells. HUVECs were stimulated with IFNγ (25 IU/mL) and TNFα (5ng/mL) for 12 h.

#### Coculture of HUVEC with JURKAT cells

After 12 h, Atg5KO HUVECs were incubated with αVCAM1 antibody (R and D #BBA5, 2.5 µg/mL) for 30 mins at 37°C. Together with αVCAM1 antibody, Invitrogen™ CellTrace™ Calcein Green (ThermoFisher Scientific # C34852) was added to HUVECs at 10nM concentration. JURKAT T cells were trypsinized and incubated with 10nM CellTrace™ Calcein Red-Orange (ThermoFisher Scientific # C34851) for 30 min. Both HUVECs and JURKATs cells were co-incubated at 1:1 ratio for 30 min at 37°C. Co-incubated cells were transferred on ice and Amnis Imagestream was used to acquire data. Ideas Application version 6.0 was used to quantify the number of doulets.

All multi color flow cytometry experiments included single color compensation/unmixing controls. Single color compensation controls or unmixing controls were prepared and run simultaneously with each experiment. For single color controls, UltraComp eBeads™ Compensation Beads (Invitrogen #01-2222-42) were stained with fluroscenly tagged antibodies individually using the same protocols as cells. Gates were defined based on FMO controls. Data analysis was perform in FlowJo V10.8.1 or FCS express 7.14 research edition.

### Immunofluorescent imaging

#### Tissues

For the preparation of frozen sections, excised tumors were fixed in 2% paraformaldehyde (PFA) at 4°C overnight, followed by 30% sucrose at 4°C overnight before being embedded in OCT (Leica). 10μm thick tumor tissue sections were stained with anti-CD3 (clone 17A2; BioLegend #100236), anti-CD31 (Abcam #28364), anti-CD31 (Millipore #MAB1398Z), anti-NG2 (Merck #AB5320), anti-aSMA (abcam #ab5694), anti-P62 (Sigma #P0067), anti-VCAM1 (Invitrogen #14-1061-82) anti-ICAM1 (BD Pharmigen #550287), anti-ICAM1 (R&D #BBA3), anti-STING (Proteintech 19851-1-AP), anti-NF-KB2 (Novus #87760), anti-IFN-B (Thermo #PA5-20390), anti-P65 (abcam #ab16502). When primary antibodies were unlabeled, fluorophore-conjugated secondary antibodies were used (Invitrogen #A-11008, #A-11006, #A78963, #A-11012, #A32733, #A78967). Images were taken with an Observer Z1 microscope (Zeiss) linked to an AxioCam MRM camera (Zeiss) with objectives of 40X or 63X magnification, Zeiss Axioscan (20x magnification, or Zeiss Airyscan for super resolution (63X). Image analysis and quantitation were performed using ImageJ software version 2.0.0.

#### Cells

For microscopy analysis, cells were grown on 1.5 mm coverslips. Fixation was performed using 4% PFA in PBS for 10 minutes, followed by permeabilization/blocking in 0.1% saponin 5% normal goat serum in PBS for 1 hour. Cells were incubated with primary antibodies overnight in a humidified chamber at 4°C followed by washing before addition of secondary antibodies for 1 h at room temperature. Nuclei were stained using DAPI. Coverslips were mounted in a drop of Prolong® Gold (Thermo Fisher Scientific #P36934). The following primary antibodies were used: anti-dsDNA (abcam #ab27156), anti-TOM20 (abcam #ab186735), anti-STING (Proteintech #19851-1-AP), anti-LMAN1 (Invitrogen #MA5-25345). Secondary antibodies used for detection were as follows: Alexa Fluor 488 goat anti-mouse (Invitrogen #A11029), Alexa Fluor 488 goat anti-rabbit (Invitrogen #A11034), Alexa Fluor 647 goat anti-mouse (Invitrogen #A21236), Alexa Fluor 647 goat anti-rabbit (Invitrogen #A21245). Super-resolution Airyscan images were acquired on an inverted Zeiss LSM880 with Airyscan microscope (Carl Zeiss). Images were collected using a 63x objective (NA 1.4). 488 and 640 nm laser lines were used, and refractive index-matched immersion oil (Zeiss) was used for all experiments. For z-stacks collection, the software-recommended optimal slice sizes were used. The setup was controlled by ZEN black (software version, Carl Zeiss Microscopy GmbH). For post processing and image analysis, Fiji/ImageJ (software version 2.9.0/1.53t) and Imaris (software version 9.1) softwares were used. The authors gratefully acknowledge the VIB Bio Imaging Core for their support & assistance in this work.

### RNA isolation

RNA from HUVECs was extracted by the recommended protocol by Qiagen RNeasy mini kit (Qiagen #74136). For muTECs, cells were sorted directly into 1mL Trizol at 4°C. Each sample was immediately frozen on dry ice and then kept at -80°C. Recommeded Trizol method was used to extract pure RNA in 12 μL nuclease free water. As suggested in the protocol, glycogen (Roche # 10901393001) was added with iso-propanol to facilitate RNA precipitation and visualization of the pellet. RNA was quantified by Agilent RNA 6000 Pico Kit (Agilent #5067-1513). RNA from blood was extracted as per the protocol recommended by the manufacturer (PureLink™ Total RNA Blood Kit #K156001).

### Quantitative RT-PCR

Specific primers were designed in Primer3 or adopted from published literature. For HUVECs, blood and tumor, RNA was converted to cDNA by QuantiTect Reverse Transcription Kit (Qiagen #205311). For muTECs, RNA was converted to cDNA by SuperScript™ III First-Strand Synthesis System (Invitrogen #18080051). ORA™ SEE qPCR Green ROX L Mix (highQu # QPD0505) was used to quantify gene expression in Applied Biosystems QuantStudio 5 Real-Time PCR System. Fold change was calculated by 2^(-ddct) method. Ct values for each gene was normalized for loading as follows. HUVEC – PPIB, blood – HPRT, muTECs – average of 18s rRNA+GAPDH. For verification of loss of Exon 3 in Atg5^BECKO^ mice, forward primer was designed to bind in Exon 3 region and reverse primer was designed to bind exon 4 reigon of *ATG5* gene.

### Western blotting

Immunoblotting on whole cell lysates was performed as previously described^26^. Membranes were incubated overnight with antibodies against VCAM1 (1:1000 dilution; Abcam #ab134047), TBK1 (1:1000 dilution; CST# 3504S), pTBK1 (1:1000 dilution; CST# 5483S), STING (1:1000 dilution; CST# 13647S), cGAS (1:1000 dilution; CST# 15102), ATG5 (1:1000 dilution; CST# 12994S), GAPDH (1:3000 dilution; CST# 5174S), VDAC (1:1000 dilution; CST# 4661S), Caspase3 (1:1000 dilution; CST# 9662S), cleaved caspase 3 (1:1000 dilution; CST# 9664S), LC3B (1:1000 dilution; CST# 3868S) and P62 (1:1000 dilution; CST# 39749S). Membranes were then incubated with respective anti-rabbit (1:2000 dilution; HRP conjugated CST# 7074S or DyLight™ 800 conjugated Thermo #SA5-10036 or DyLight™ 680 conjugated Thermo #35568) or anti-mouse secondary antibodies (1:2000 dilution; HRP conjugated CST# 7076S) for 1h. Protein bands were detected either using Amersham ECL imager (for HRP conjugated antibodies) or Amersham Typhoon Biomolecular Imager (for DyLight conjufgated antibodies). BIO-RAD Clarity and Clarity Max ECL Western Blotting Substrates were used with HRP conjugated antibodies. For semi-native western blotting for STING protein, same protocol was following with an exception of non-denaturing conditions while preparing cell lysates. Quantification was done using Image J (NIH).

### Nanostring

nCounter PanCancer Mouse Immune Profiling Panel (NanoString #XT-CSO-MIP1-12) was used to screen gene expression in muTEC. Data was analyzed with nSolver software version 4.0 (advanced analysis). Ten ng RNA from muTECs was loaded for analysis. *GSEA analysis*: Gene set enrichment analysis (GSEA) was carried out using GSEA software with MH: hallmark gene sets in the MSigDB database on the list of 744 genes analysed as input through Nanostring technology. The default gene list was used as background. For visualization, SRplot online tool was used to build Bubble plot, in which p values are represented by colors, gene counts are represented by bubble size.

### Single cell RNA sequencing analysis – metadata analysis

#### Patient population

We collected 79 fresh tissue samples from 53 patients across 4 tumor types: breast cancer (BC; n=31, 39%), colorectal cancer (CRC; n=21, 27%), high-grade serous ovarian carcinoma (HGSOC; n=12, 15%) and non-small cell lung cancer (NSCLC; n=15, 19%). All samples were treatment-naïve tumors that have already been published^88,89^. The local ethics committee at the University Hospital Leuven approved the single-cell study for each cancer type, and all patients provided written informed consent.

### Single cell RNA sequencing (scRNA-SEQ) and expression analysis

Single-cell suspensions underwent 3’ or 5’ scRNA-seq using the ChromiumTM Single Cell V(D)J Solution from 10x Genomics as previously described^89^. Most tissues (CRC, HGSOC and NSCLC samples; 66%) were subjected to 3’ scRNAseq, although in some cases (BC samples; 34%) 5’ were performed instead. Gene expression libraries were generated according to the manufacturer’s instructions with the final aim to obtain 5,000 cells per sample. All the libraries were sequenced on Illumina NextSeq, HiSeq4000 and/or NovaSeq6000 and mapped to the GRCh38 human reference genome using CellRanger (10x Genomics). The latter was also used to generate raw gene expression matrices19, then analyzed using the Seurat v4 R package20. Cells expressing <200 or >6000 genes, <400 unique molecular identifiers (UMIs) and >25% mitochondrial counts were removed.

### Clustering ECs subtypes

We first analyzed each cancer type independently and focused on the clusters analysis as previously described^88,89.^ All cells assigned as ECs per each cancer type were merged and the same clustering strategy was applied at the subclusters level. In addition, for the identification of cellular subgroups we integrated the data generated from different technologies (3’ or 5’ scRNA-seq) with the batch effect correction algorithm Harmony (Korsunsky, I. et al, Nat. Methods 2019). In addition to the previously mentioned confounding factors, at the subclusters level we also regressed out for individual tumor types, interferon (IFN) response and stress signature, as previously described^88^. Clusters representing distinct cell types were identified based on the expression of marker genes.

### Statistical analysis

Statistical analyses were performed using R (version 4.0.3, R Foundation for Statistical Computing, R Core Team, Vienna, Austria). Spearman’s correlation analysis was applied to quantify correlations between levels of signatures gene expression. Statistical analyses were performed with the Mann-Whitney test, using a two-sided alternative hypothesis at the 5% significance level.

### Melanoma single cell EC data

The raw count matrix of the entire Grand Challenge melanoma dataset was obtained and anaysed as in described in the pre-print publication (https://doi.org/10.1101/2022.08.11.502598). Tumour microenvirontment cells including the CD8+ Tcells were identified as in Landeloos et. al. (unpublished) and the correlations of CD8+ Tcells with muTEC-DE score were performed using Spearman test. Endothelial cells were identified by calculating the AUCell gene enrichment score of EC signatures and selecting cells with an AUCell score > 0.1 for downstream analysis. High variable genes were selected using the FindVariableFeatures function and auto-scaled with the ScaleData function. Principal component analysis (PCA) was performed using the default RunPCA function in the Seurat package^90^, and batch effect correction was applied to each sample using the Harmony algorithm^91^ based on PCA space. The data were visualized using uniform manifold approximation and projection (UMAP) with the RunUMAP step (dims = 10), and unsupervised clustering was performed using the FindNeighbors followed by FindClusters function (dims = 10, resolution = 0.4) in the Seurat package. EC subtypes were identified primarily based on marker genes reported in the literature^92^. The enrichment of given gene sets for each cell was evaluated using the AUCell package^93^ for Gene Set Enrichment Analysis (GSEA). Modified violin + box plots were generated using customized R code, and these functions were integrated into the R package “SeuratExtend,” which is available on Github (https://github.com/huayc09/SeuratExtend). Statistical analyses were performed using ggpubr package in R.

### Clinical data for MILAN analysis

A selection of patients included in a previously published data set ^94^ was made to be analyzed with the MILAN technology. 12 pretreatment, formalin-fixed, paraffin-embedded (FFPE) melanoma metastasis samples from 12 patients were collected from the histopathological archives of the University Hospital of Leuven (Clinical data of the selected patient are reported in Extended data Table 2). All patients were treated with anti-PD-1 monotherapy (nivolumab or pembrolizumab) and after the biopsy was taken. Only biopsies taken < 365 days before the start of anti-PD-1 monotherapy were included. Furthermore, only patients with measurable disease were selected, hence enabling tumor response assessment according to RECIST 1.1^95^. Patients were classified according to the best objective response to immunotherapy during their time of follow-up, as defined by RECIST 1.1^95^. Complete response and partial response were classified as R for responder, progressive disease or stable disease as NR for non responder. According to these criteria, 6 patients could be classified as R (six samples) and 6 patients as NR (six samples). Only metastatic samples were eligible for inclusion. An expert dermatopathologists specialized in melanoma research (FMB) selected the most representative areas of the tumors for tissue microarray (TMA) construction. For each metastasis, one to five representative cores/regions of interest were sampled having at least a size of 1 mm in diameter. The number of samples taken was determined by the specimen and the morphologic heterogeneity of both the melanoma and the inflammatory infiltrate. Therefore, a smaller number of cores were taken from small and homogeneous samples whereas a larger number was taken from large but heterogeneous specimens. The studies were conducted in accordance with recognized ethical guidelines (Declaration of Helsinki). This project was approved by the Ethical Commission of the University Hospital of Leuven and approved by the review board.

### Multiple Iterative Labeling by Antibody Neodeposition (MILAN)

*Tissue staining:* Multiplex immunofluorescent staining was conducted using the MILAN protocol as described previously^96^, Immunofluorescence images were acquired using the Axio scan.Z1 slidescanner (Zeiss, Germany) at 10X objective with a resolution of 0.65 micron/pixel at 16 bit color depth. Hematoxylin and eosin (H&E) slides were also digitized using the same slidescanner in brightfield modus with a 10X objective at 8 bit color depth. All samples were stained together, and the quality of the stains was evaluated by an experienced pathologist (FMB). Poor quality areas showing artifacts such as tissue folds or antibody aggregation were excluded from downstream analysis.

### Image analysis

Image analysis was conducted using a previously described^94^ customized pipeline. Briefly, images were corrected for field of view artifact using a method described in the literature^97^, Then, the overlapping regions of adjacent tiles were stitched together by minimizing the Frobenius distance. Next, images from consecutive rounds were aligned (registered) following an algorithm previously described^98^. To register the images from consecutive rounds, the first round was set as a fixed image while all the following rounds were used as moving images. The DAPI channel was used to calculate transformation matrices, which were then applied to the other channels. The quality of the overlapping regions was visually evaluated and poor registered areas were removed from downstream analyses. Then, tissue autofluorescence was removed by subtracting a baseline image with only secondary antibody. Finally, cell segmentation was applied to the DAPI channel using STARDIST^99,100^. which delineates a contour for each cell present in the tissue. For each of these cells, the following features were extracted: topological features (X/Y coordinates), morphological features (nuclear size), and molecular features (Mean Fluorescence Intensity (MFI) of each measured marker).

### Single-cell analysis

MFI values were normalized using Z-scores within each sample, as recommended in Caicedo et al^101^. To avoid a strong influence from outliers in downstream analyses, Z-scores were trimmed within the [-5, 5] range. Three different clustering methods were used to map single cells to known cell phenotypes: PhenoGraph^102^, FlowSom^103^, and KMeans, which were implemented in the Rphenograph, FlowSOM, and stats R packages, respectively. While FlowSom and KMeans required the number of clusters as input, PhenoGraph could be executed by exclusively defining the number of nearest neighbors to calculate the Jaccard coefficient 20 (standard value). The number of clusters identified by PhenoGraph was then used as an argument for FlowSom and KMeans. Clustering was performed exclusively on a subset of the identified cells (2.500), which were selected by stratified proportional random sampling and using the following markers: AQP1, CD31, CD4, CD8, MelanA, and SOX10. For each clustering method, clusters were mapped to known cell phenotypes based on manual annotation by domain experts. For every cell, if two or more clustering methods agreed on the assigned phenotype, the cell was annotated accordingly. However, if all three clustering methods disagreed on the assigned phenotype, the cell was annotated as “not otherwise specified” (NOS).

Following consensus clustering, 4 different cell types were identified (+NOS): Melanoma (SOX10+ | MelanA+), Blood Vessels (AQP1+ | CD31+), T-helpers (CD4+), and Cytotoxic T cells (CD8+). To extrapolate the cell labels to the remaining cells in the dataset, a uMap was built by sampling 500 cells for each identified cell type in the consensus clustering, and the entire dataset was projected into the uMap using the base predict R function. For each cell, the label of the closest 100 neighbors was evaluated in the uMap space, and the most frequent cell type was assigned as the label. Digital reconstructions of the tissue samples were obtained by coloring the segmentation mask by the assigned cell label. An experienced dermatopathologist (FMB) used these reconstructions to annotate different areas of interest: “tumoral bulk” areas, tumor-stroma interface, and non-tumoral areas. Relative cytometry enrichment was performed for each individual area using Wilcoxon rank-sum tests. Given the exploratory nature of the analysis, p-values were not adjusted for multiple comparisons. Statistical analysis and data presentation were performed using R Studio (version RStudio 2022.07.2). Relative STING+ and VCAM1+ blood vessels were identified by applying a cutoff on the respective normalized data. The same analysis was performed for Cytotoxic T cells using PD-1 and GrzB.

### Statistical Analysis

Statistical difference between two groups with equal sample size was determined by standard two-sided Student’s t-test. For comparing two groups with unequal sample size, Mann–Whitney U test was used. To compare paired samples from invitro experiments, ratio paried t-test was used. For comparing three or more groups, one-way ANOVA with Tukey’s post-hoc test or Holm-Sidak correction for multiple comparisons were used. Statistical outliers were excluded based on the on Grubb’s test in Prism software. All TEC/tissue analysis was conducted with at least three mice per group or HUVECs with three independent donors. GP p value style was used. */$/represents a P-value < 0.05, **/$$ < 0.01, *** p < 0.001 and **** p < 0.0001 where a p-value < 0.05 was used to determine statistical difference. All statistical analysis was performed in Prism 9.4.1.

## Acknowledgments

We thank E. Vervoot, K. Rillaerts, B. Gilbert for excellent expert technical support. We thank Theo Killian for initial bioinformatics support. The authors gratefully acknowledge the VIB Bio Imaging Core and Flow Core for their support & assistance in this work. P.A. is supported by grants from the Flemish Research Foundation (FWO-Vlaanderen; G076617N, G049817N, G070115N), the EOS MetaNiche consortium N° 40007532, Stichting tegen Kanker (FAF-F/2018/1252) and the iBOF/21/053 ATLANTIS consortium with G.B and M.JM.B. M.JM.B is supported by grants from FWO-Vlaanderen (G035320N, G044518N, EOS CD-INFLADIS) and Ghent University (BOF23/GOA/001). J.V.,F.R. and D.H. received FWO Doctoral Fellowships and K.J. of a Postdoctoral fellowship (12Y4322N) from the Flemish Research Foundation (FWO-Vlaanderen), Belgium.

