## Extended data figures for "TUMOR ENDOTHELIAL CELL AUTOPHAGY IS A KEY VASCULAR-IMMUNE CHECKPOINT IN MELANOMA"

**Extended data Fig 1. Validation of genetic loss of *Atg5* in tumor endothelial cells**

**a**, gating strategy for quantifying GFP signal in $CD31^+$ tumor endothelial cells in subcutaneous B16-F10 tumors from WT and $Atg5^{BECKO}$ mice with subcutaneous B16-F10 tumors. **b**, Representative immunofluorescence images showing tdtomato and GFP in $CD31^+$ tumor endothelial cells in subcutaneous B16-F10 tumors from WT and $Atg5^{BECKO}$ mice with subcutaneous B16-F10 tumors. **c**, Gene expression of *Atg5* normalized to 18s rRNA+GAPDH in sorted $CD31^+$ tumor endothelial cells in subcutaneous B16-F10 tumors from WT and $Atg5^{BECKO}$ mice. Forward primer was designed to bind in Exon 3 region and reverse primer was designed to bind in Exon 4 region of *Atg5* gene. **d-f**, Representative immunofluorescence images and quantification for p62 (MFI) **(d)**, NG2 (% coveragel**(e)** and αSMA (% coverage) **(f)** in CD31+ tumor endothelial cells from subcutaneous B16-F10 tumors from WT and $Atg5^{BECKO}$ mice. Scale bars represent 10μm. For immunofluorescence staining, at least 3 WT and 3 $Atg5^{BECKO}$ mice were used for the quantification. All data represent mean ± sem. Statistical differences were determined using two-sided Student's t-test. Source data are available for **a-f**.

**Extended data Fig 2. Immunophenotyping of WT and $Atg5^{BECKO}$ mice with subcutaneous B16-F10 melanoma.**

**a,** Gating strategy for analyzing immune cell subsets in subcutaneous B16-F10 melanoma from WT and $Atg5^{BECKO}$ Mice. **b,** Gene expression of *CD8*, *CD3g*, *CD45* and *Nkp46* in blood collected from WT and $Atg5^{BECKO}$ mice with subcutaneous B16-F10 tumors and injected with αCD8 antibody. All data represent mean ± sem. Statistical differences were determined using one-way Anova with Tukey corrections for multiple comparisons **(b).** Source data available for **a-b**.

**Extended data Fig 3. Phenotype of TEC from WT and $Atg5^{BECKO}$ mice and human TEC subclusters from single-cell RNA-seq atlases**

**a,** flow cytometry analysis for the surface expression of MHCI and MHCII in $CD31^+$ tumor endothelial cells derived in subcutaneous B16-F10 tumor from WT and $Atg5^{BECKO}$ Mice. **b,** representative images and quantification (MFI) of immunofluorescence staining for VCAM1, ICAM1 and STING in CD31+ tumor endothelial cells from tumor sections of subcutaneous YUMMER 1.7 tumors from WT and $Atg12^{ECKO}$ mice. Scale bars represent 10μm VCAM1 and STING and 20 μm for IcAM1. **c-f,** preliminary analysis of publicly available single-cell RNA-seq atlases from primary tumors of treatment-naïve patients. Genetic markers used to cluster EC subsets and their corresponding expression shown in a heatmap **c,** Heatmap showing expression of representative marker genes across 9 ECs subtypes. **d,**

UMAP map of ECs (n= 7.573) color-coded for the indicated cell type.**e,** Pie chart showing the pan-cancer relative abundance of the 9 ECs subtypes. **f,** Relative abundance of the 9 ECs subtypes across tumor types included in the study. PCV = Post-capillary venules. HGSOC, high-grade serous ovarian carcinoma, CRC, colorectal cancer, BC, breast cancer. NSCLC, non-small cell lung cancer.For immunofluorescence staining, 2 WT and 3 Atg12$^{ECKO}$ mice were used for the analysis. All data represent mean ± SEM. Statistical differences were determined using two-sided Student's t-test **(a,b)** or In **f**, exact P values by two-sided Mann-Whitney test or two-sided Wilcoxon matched-pairs signed rank test: *P<0.05, **P<0.01, ***P<0.001. Source data are available for **a-b**.

**Extended data Fig 4. Autophagy blockade in HUVECs promotes formation of STING dimers and oligomers and accumulation in ERGIC**

**a,b,c,** representative western blot for Atg5 **(a)**, LC3B **(b)** and p62 **(c)** and quantification for p62 in HUVECs nucleofected with scrambled (Ctr) or Atg5 specific guide RNA (Atg5KO) conjugated with cas9 protein. **d,** western blot of BNIP3L, VDAC, caspase-3 and cleaved caspase 3 in Ctr and Atg5KO HUvECs. **e,** gene expression analysis of *VCAM1*, *SELE*, *ICAM1*, *CXCL10* and *C3CXL1* in HUVECs treated with vehicle (Ctr) or ULK1/2 inhibitor (top) and vehicle (Ctr) or bafilomycin A (BfA) (bottom). **f,** gating strategy for flow cytometic analysis of surface expression of VCAM1 (% of live cells) in Ctr and Atg5KO HUVECs upon stimulation with IFNγ for 4h. (top) Imaging flow cytometry analysis for doublet formation (# doublets/10,000 events) in Ctr and Atg5KO HUVECs (stimulated with IFNγ and TNFα for 24h) with JURKAT cells (bottom). **g,** representative western blot and quantification of STING monomers, dimers and oligomers in HUVECs treated with vehicle or ULK1/2 inhibitor. **h,** super-resolution Airyscan immunofluorescence images for STING (green) and LMAN1 (magenta) proteins in Ctr and Atg5KO HUVEC. Scale bars represent 10μm and at least 30 cells (10 per donor) were imaged per condilion. **i,** Super-resolution Airyscan immunofluorescence images for DAPI in Ctr and Atg5 KO HUVECs. Scale bars represent 10μm. Representative images from three independent experiments. Statistical differences were determined using two-sided Student's t-test. Source data are available for **a-i**.

**Extended data Fig 5. Analysis of genetic deletion of *Sting* and *Atg5* in TECs**

**a-b,** gene expression for *ATG5* (a) and *STING* (b) in CD31$^{+}$ tumor endothelial cells sorted from subcutaneous B16-F10 melanoma tumors from WT, Atg5$^{BECKO}$ and Atg5/STING$^{BECDKO}$ mice. For *ATG5*, forward primer was designed to bind in Exon 3 region and reverse primer was designed to bind in

Exon 4 region. For STING, forward primer designed to bind in Exon 2 region and reverse primer was designed to bind in Exon 3 region. **c,** Quantification of immunofluorescence staining for CD3+ T cells in the tumor sections of subcutaneous B16-F10 tumors from Atg5+/+, Atg5/Sting+/+, Atg5$^{BECKO}$ and Atg5/Sting$^{BECDKO}$ mice. **d**, representative images and quantification (MFI) of immunofluorescence staining for CD31 and NIK in the tumor sections of subcutaneous B16-F10 tumors from Atg5+/+, Atg5/Sting+/+, Atg5$^{BECKO}$ and Atg5/Sting$^{BECDKO}$ mice. For immunofluorescence staining, at least 3 mice per group were used for the quantification. All data show mean ± sem. using one-way Anova with Tukey corrections for multiple comparisons. Source data available for **a-c**.

**Extended data Fig 6. Interferon subtype of ECs from patients responding to αPD1 therapy upregulate muTEC-DE genes upon treatment**

**a,** grouped tumor volume of B16-F10 subcutaneous tumor bearing WT and Atg5$^{BECKO}$ mice injected with isotype (ISO) or αPD1 antibody. Data is from 1 representative experiment. **b,** UMAP map of ECs (n= 7.573) color-coded for the indicated cell type. UMAP of ECs color coded for the indicated cell types. **c,** Heatmap showing expression of representative marker genes across 11 ECs subtypes. **d,** gene enrichment score of muTEC-DE geneset across different subsets of huTECs from treatment naïve stage III/IV melanoma patients receiving anti-PD1 based therapy monotherapy (nivolumab). Median-quantile-min/max + population distribution is shown in violin + boxplot (**d**). Wilcoxon test, $*p < 0.05$; $**p < 0.01$; $***p < 0.001$; $****p < 0.0001$.

**Extended data Fig 7. Protein markers used to annotate cells in the MILAN analysis**

**a,** table illustrating the markers used for annotating ECs and CD8$^+$ T cells in tissue sections from melanoma patients undergoing anti-PD1 monotherapy using MiLAN. **b,** AQP1, CD31, CD4, CD8, MelanA, and SOX10 expression projected on the UMAP generated built by sampling 500 cells for each identified cell type in the consensus clustering. Entire dataset was projected into the uMap using the base predict R function. **c,** UMAP showing identified cell types based on the markers shown in extended figure panel a.

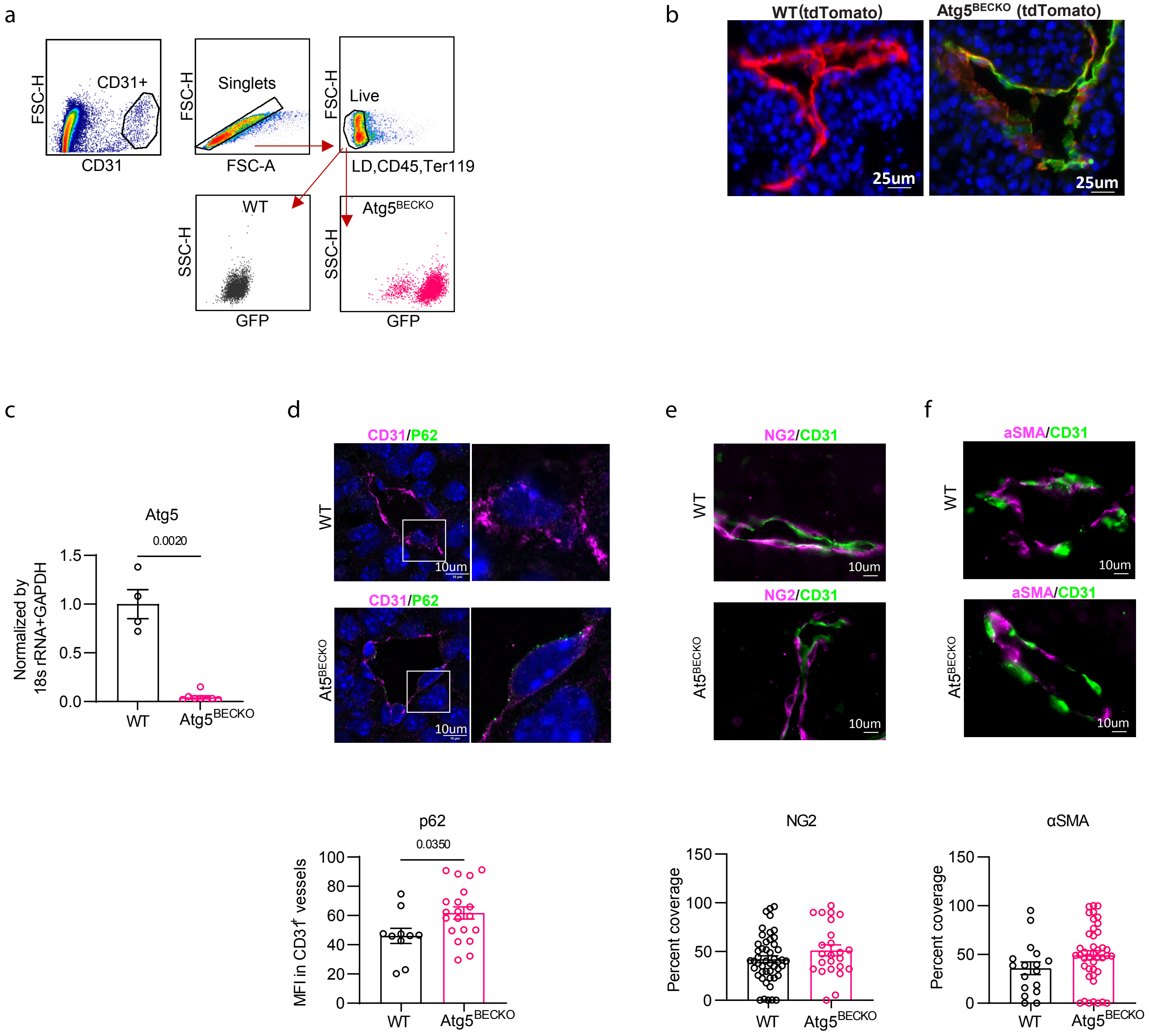
a
CD31+
FSC-H
CD31
Singlets
FSC-H
FSC-A
Live
FSC-H
LD,CD45,Ter119
WT
SSC-H
GFP
Atg5BECKO
SSC-H
GFP
b
WT(tdTomato)
Atg5BECKO (tdTomato)
25um
25um
c
Atg5
0.0020
Normalized by 18s rRNA+GAPDH
1.5
1.0
0.5
0.0
WT
Atg5BECKO
d
CD31/P62
WT
10um
CD31/P62
At5BECKO
10um
e
NG2/CD31
WT
10um
NG2/CD31
At5BECKO
10um
f
aSMA/CD31
WT
10um
aSMA/CD31
At5BECKO
10um
p62
0.0350
MFI in CD31+ vessels
100
80
60
40
20
0
WT
Atg5BECKO
NG2
Percent coverage
150
100
50
0
WT
Atg5BECKO
αSMA
Percent coverage
150
100
50
0
WT
Atg5BECKO

a

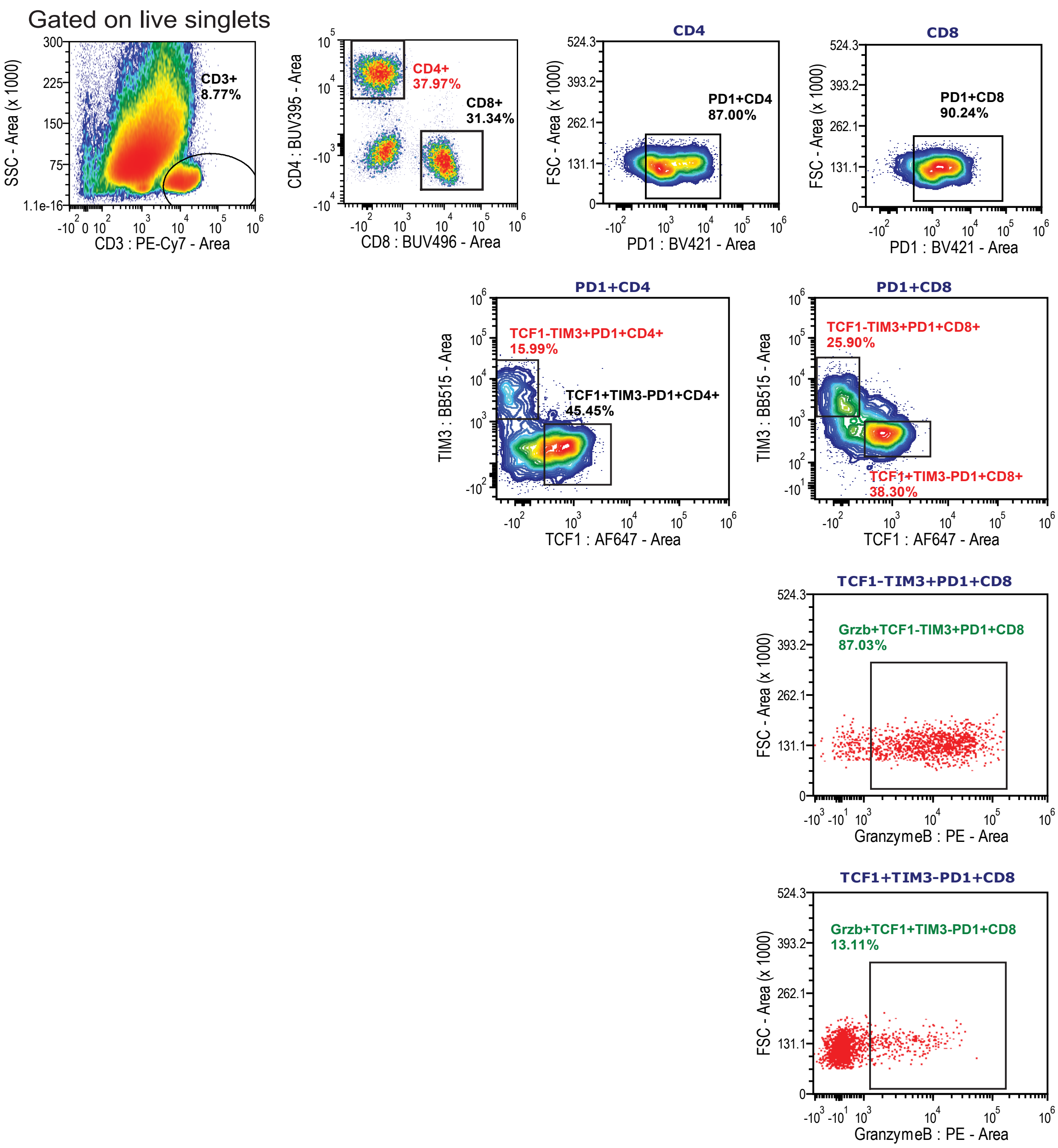
Gated on live singlets
CD3+
8.77%
SSC - Area (x 1000)
CD3 : PE-Cy7 - Area
CD4+
37.97%
CD8+
31.34%
CD4 : BUV395 - Area
CD8 : BUV496 - Area
CD4
PD1+CD4
87.00%
FSC - Area (x 1000)
PD1 : BV421 - Area
CD8
PD1+CD8
90.24%
PD1+CD4
TCF1-TIM3+PD1+CD4+
15.99%
TCF1+TIM3-PD1+CD4+
45.45%
TIM3 : BB515 - Area
TCF1 : AF647 - Area
PD1+CD8
TCF1-TIM3+PD1+CD8+
25.90%
TCF1+TIM3-PD1+CD8+
38.30%
TCF1-TIM3+PD1+CD8
Grzb+TCF1-TIM3+PD1+CD8
87.03%
GranzymeB : PE - Area
TCF1+TIM3-PD1+CD8
Grzb+TCF1+TIM3-PD1+CD8
13.11%

b

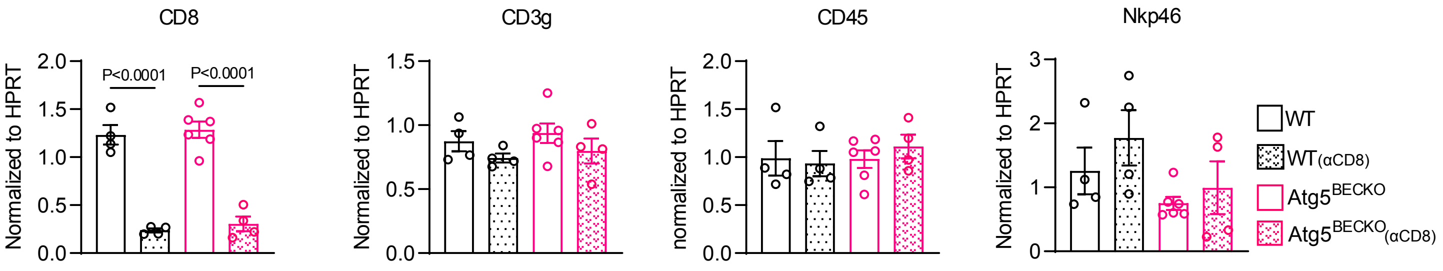
CD8
CD3g
CD45
Nkp46
Normalized to HPRT
normalized to HPRT
P<0.0001
P<0.0001
WT
WT(αCD8)
Atg5BECKO
Atg5BECKO(αCD8)

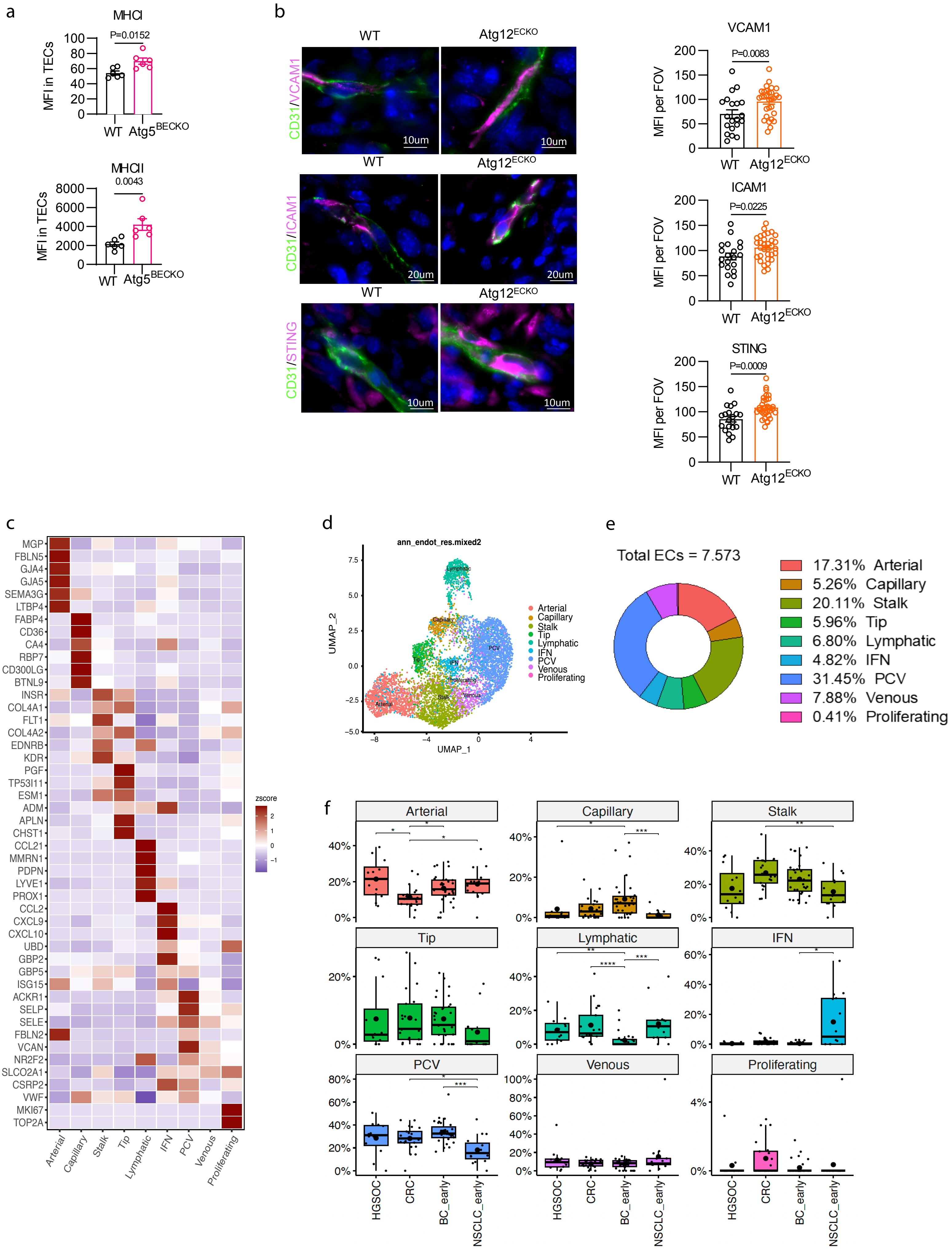
a
MHCI
P=0.0152
MFI in TECs
100
80
60
40
20
0
WT
Atg5BECKO
MHCII
0.0043
MFI in TECs
8000
6000
4000
2000
0
WT
Atg5BECKO
b
WT
Atg12ECKO
CD31/VCAM1
10um
CD31/ICAM1
20um
CD31/STING
10um
VCAM1
P=0.0083
MFI per FOV
200
150
100
50
0
WT
Atg12ECKO
ICAM1
P=0.0225
MFI per FOV
STING
P=0.0009
MFI per FOV
c
MGP
FBLN5
GJA4
GJA5
SEMA3G
LTBP4
FABP4
CD36
CA4
RBP7
CD300LG
BTNL9
INSR
COL4A1
FLT1
COL4A2
EDNRB
KDR
PGF
TP53I11
ESM1
ADM
APLN
CHST1
CCL21
MMRN1
PDPN
LYVE1
PROX1
CCL2
CXCL9
CXCL10
UBD
GBP2
GBP5
ISG15
ACKR1
SELP
SELE
FBLN2
VCAN
NR2F2
SLCO2A1
CSRP2
VWF
MKI67
TOP2A
zscore
2
1
0
−1
Arterial
Capillary
Stalk
Tip
Lymphatic
IFN
PCV
Venous
Proliferating
d
ann_endot_res.mixed2
UMAP_2
UMAP_1
7.5
5.0
2.5
0.0
−2.5
−5.0
−8
−4
0
4
Arterial
Capillary
Stalk
Tip
Lymphatic
IFN
PCV
Venous
Proliferating
e
Total ECs = 7.573
17.31% Arterial
5.26% Capillary
20.11% Stalk
5.96% Tip
6.80% Lymphatic
4.82% IFN
31.45% PCV
7.88% Venous
0.41% Proliferating
f
Arterial
Capillary
Stalk
Tip
Lymphatic
IFN
PCV
Venous
Proliferating
HGSOC
CRC
BC_early
NSCLC_early
0%
20%
40%
60%
80%
100%
4%

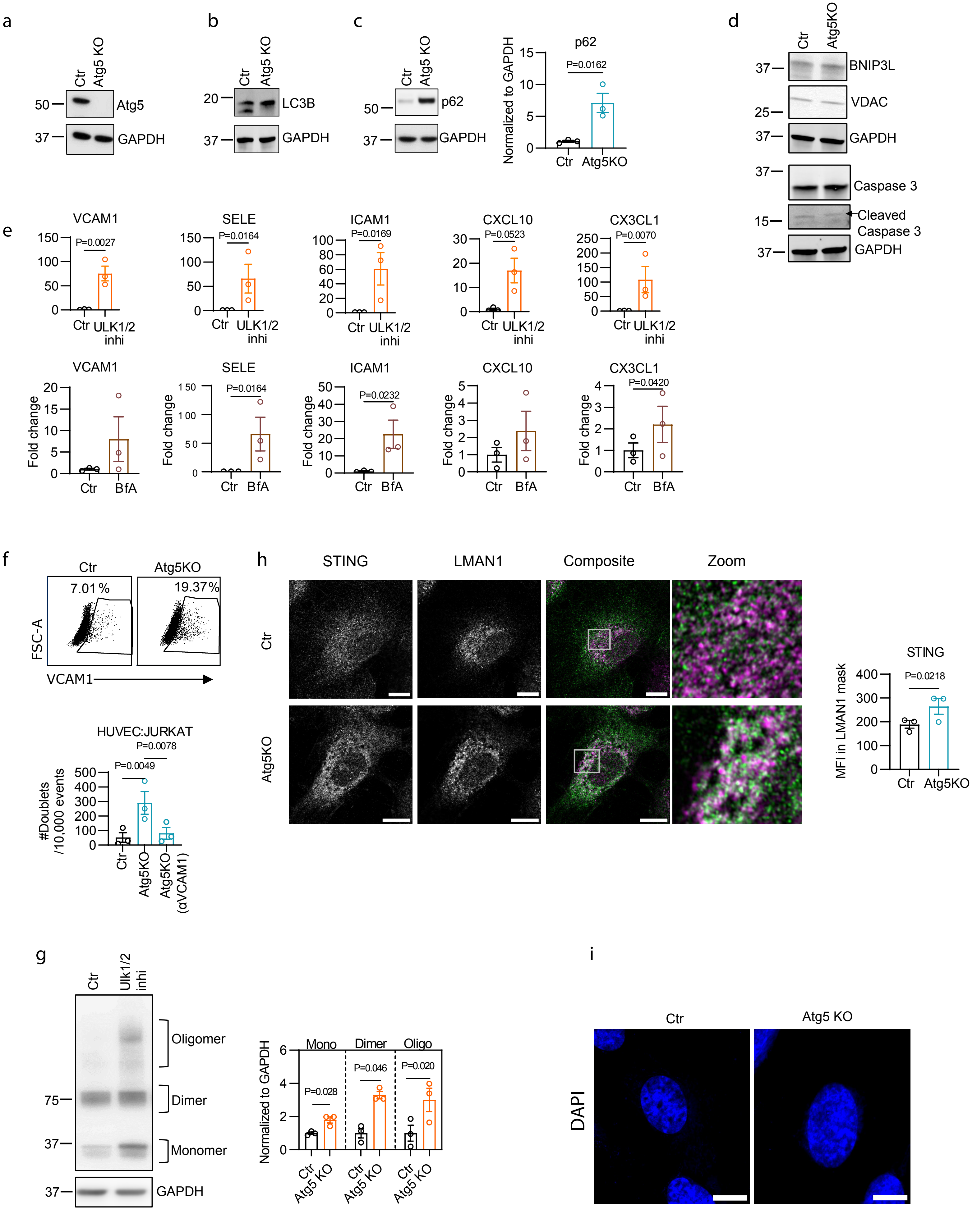
a
Ctr
Atg5 KO
50
Atg5
37
GAPDH
b
Ctr
Atg5 KO
20
LC3B
37
GAPDH
c
Ctr
Atg5 KO
50
p62
37
GAPDH
p62
P=0.0162
Normalized to GAPDH
Ctr
Atg5KO
d
Ctr
Atg5KO
37
BNIP3L
VDAC
25
37
GAPDH
37
Caspase 3
15
Cleaved
Caspase 3
37
GAPDH
e
VCAM1
P=0.0027
SELE
P=0.0164
ICAM1
P=0.0169
CXCL10
P=0.0523
CX3CL1
P=0.0070
Ctr
ULK1/2
inhi
VCAM1
SELE
P=0.0164
ICAM1
P=0.0232
CXCL10
CX3CL1
P=0.0420
Fold change
Ctr
BfA
f
Ctr
Atg5KO
7.01%
19.37%
FSC-A
VCAM1
HUVEC:JURKAT
P=0.0078
P=0.0049
#Doublets
/10,000 events
Ctr
Atg5KO
Atg5KO
(αVCAM1)
h
STING
LMAN1
Composite
Zoom
Ctr
Atg5KO
STING
P=0.0218
MFI in LMAN1 mask
Ctr
Atg5KO
g
Ctr
Ulk1/2
inhi
Oligomer
75
Dimer
37
Monomer
37
GAPDH
Mono
Dimer
Oligo
P=0.028
P=0.046
P=0.020
Normalized to GAPDH
Ctr
Atg5 KO
i
Ctr
Atg5 KO
DAPI

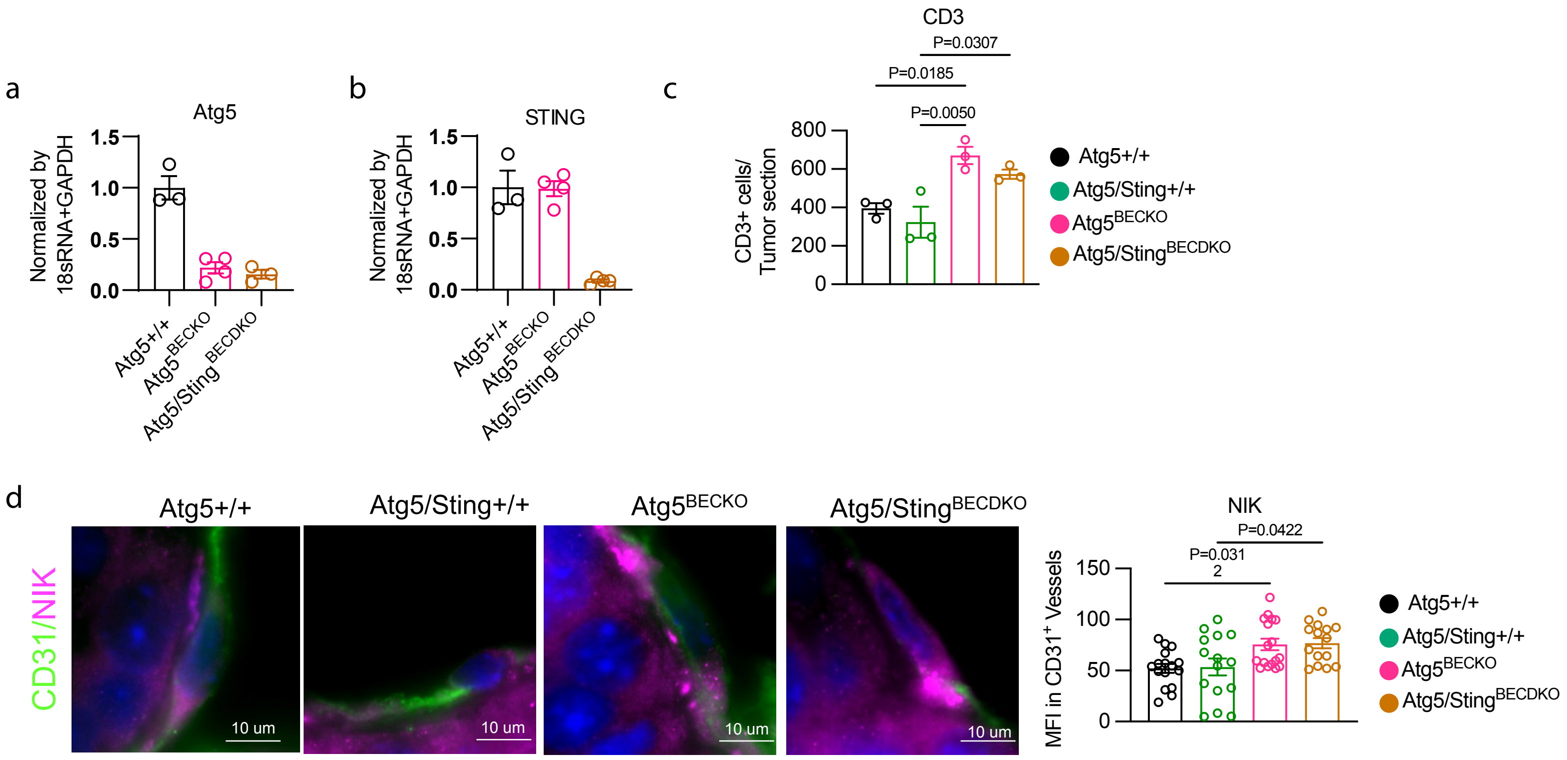

a
Atg5
Normalized by 18sRNA+GAPDH
1.5
1.0
0.5
0.0
Atg5+/+
Atg5BECKO
Atg5/StingBECDKO
b
STING
Normalized by 18sRNA+GAPDH
1.5
1.0
0.5
0.0
Atg5+/+
Atg5BECKO
Atg5/StingBECDKO
c
CD3
P=0.0307
P=0.0185
P=0.0050
CD3+ cells/ Tumor section
800
600
400
200
0
Atg5+/+
Atg5/Sting+/+
Atg5BECKO
Atg5/StingBECDKO
d
Atg5+/+
Atg5/Sting+/+
Atg5BECKO
Atg5/StingBECDKO
CD31/NIK
10 um
10 um
10 um
10 um
NIK
P=0.0422
P=0.031 2
MFI in CD31+ Vessels
150
100
50
0
Atg5+/+
Atg5/Sting+/+
Atg5BECKO
Atg5/StingBECDKO

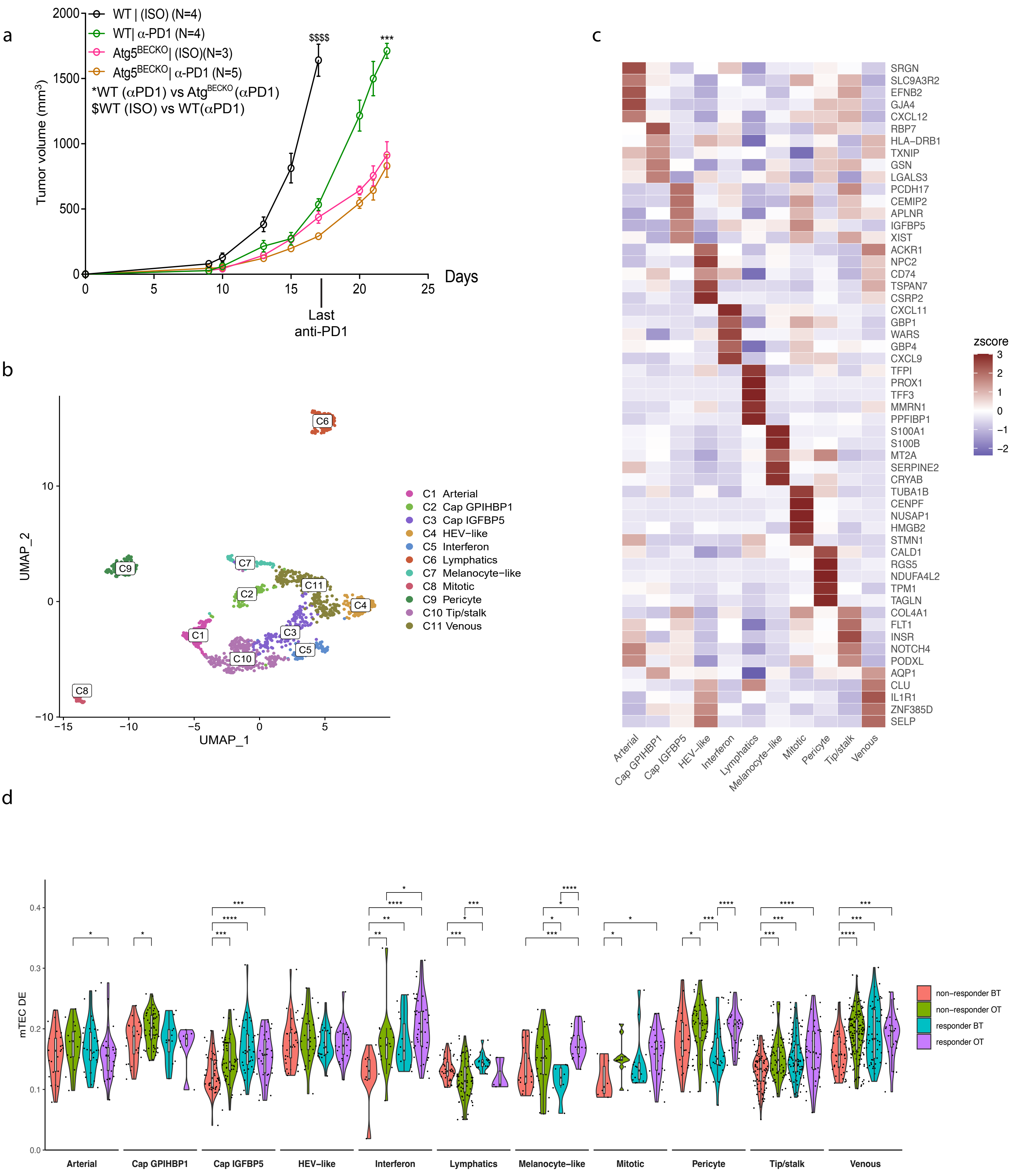
a
WT | (ISO) (N=4)
WT| α-PD1 (N=4)
Atg5BECKO| (ISO)(N=3)
Atg5BECKO| α-PD1 (N=5)
*WT (αPD1) vs AtgBECKO (αPD1)
$WT (ISO) vs WT(αPD1)
$$$$
***
Tumor volume (mm3)
2000
1500
1000
500
0
0
5
10
15
20
25
Days
Last
anti-PD1
b
UMAP_2
UMAP_1
C1 Arterial
C2 Cap GPIHBP1
C3 Cap IGFBP5
C4 HEV−like
C5 Interferon
C6 Lymphatics
C7 Melanocyte−like
C8 Mitotic
C9 Pericyte
C10 Tip/stalk
C11 Venous
c
zscore
SRGN
SLC9A3R2
EFNB2
GJA4
CXCL12
RBP7
HLA-DRB1
TXNIP
GSN
LGALS3
PCDH17
CEMIP2
APLNR
IGFBP5
XIST
ACKR1
NPC2
CD74
TSPAN7
CSRP2
CXCL11
GBP1
WARS
GBP4
CXCL9
TFPI
PROX1
TFF3
MMRN1
PPFIBP1
S100A1
S100B
MT2A
SERPINE2
CRYAB
TUBA1B
CENPF
NUSAP1
HMGB2
STMN1
CALD1
RGS5
NDUFA4L2
TPM1
TAGLN
COL4A1
FLT1
INSR
NOTCH4
PODXL
AQP1
CLU
IL1R1
ZNF385D
SELP
Arterial
Cap GPIHBP1
Cap IGFBP5
HEV-like
Interferon
Lymphatics
Melanocyte-like
Mitotic
Pericyte
Tip/stalk
Venous
d
mTEC DE
non−responder BT
non−responder OT
responder BT
responder OT
Arterial
Cap GPIHBP1
Cap IGFBP5
HEV−like
Interferon
Lymphatics
Melanocyte−like
Mitotic
Pericyte
Tip/stalk
Venous

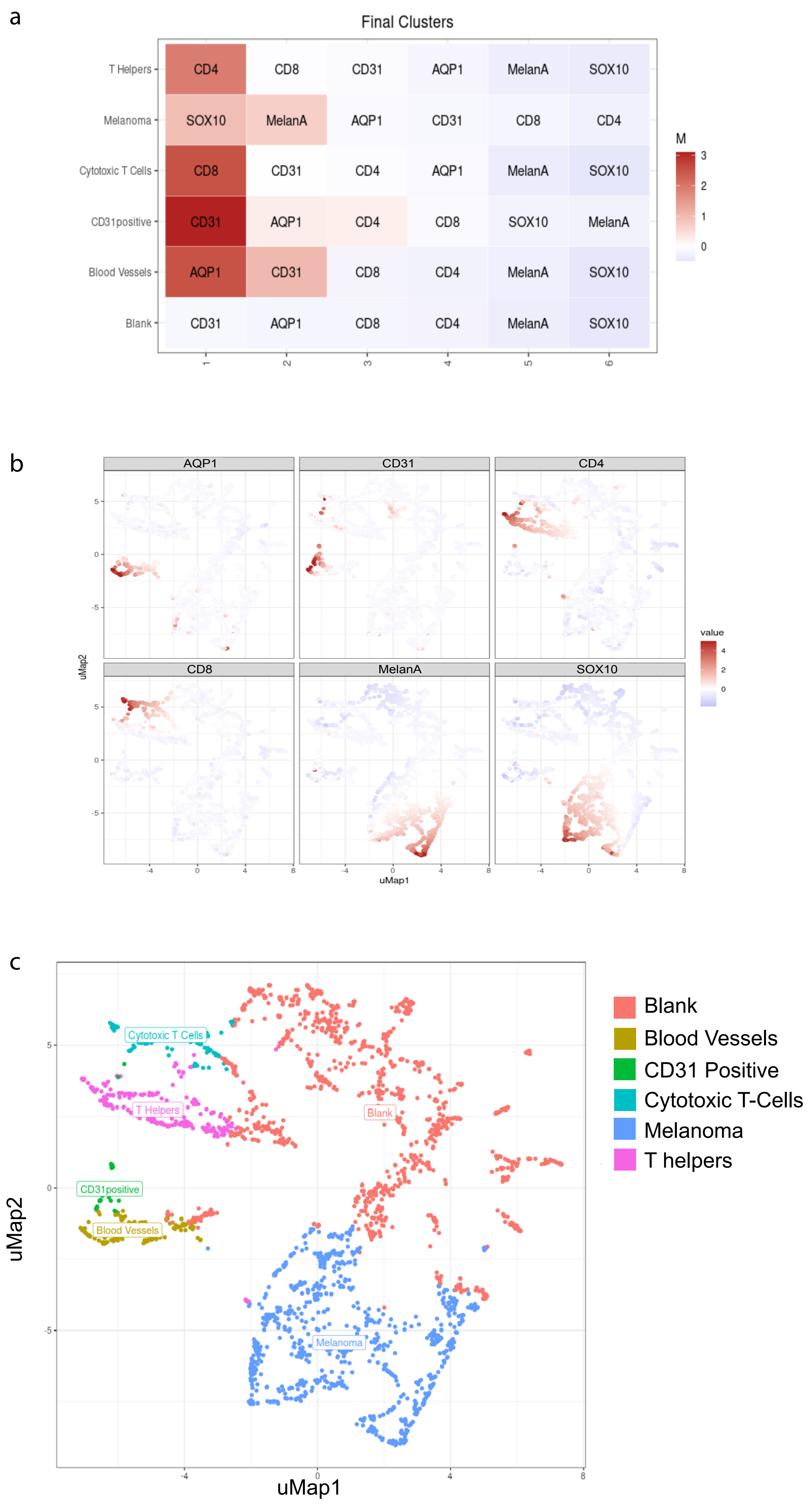

a
Final Clusters
T Helpers: CD4 CD8 CD31 AQP1 MelanA SOX10
Melanoma: SOX10 MelanA AQP1 CD31 CD8 CD4
Cytotoxic T Cells: CD8 CD31 CD4 AQP1 MelanA SOX10
CD31positive: CD31 AQP1 CD4 CD8 SOX10 MelanA
Blood Vessels: AQP1 CD31 CD8 CD4 MelanA SOX10
Blank: CD31 AQP1 CD8 CD4 MelanA SOX10
1 2 3 4 5 6
M
3 2 1 0
b
AQP1
CD31
CD4
CD8
MelanA
SOX10
uMap2
uMap1
value
4 2 0
5 0 -5
-4 0 4 8
c
Cytotoxic T Cells
T Helpers
Blank
CD31positive
Blood Vessels
Melanoma
uMap2
uMap1
5 0 -5
-4 0 4 8
Blank
Blood Vessels
CD31 Positive
Cytotoxic T-Cells
Melanoma
T helpers
